# DNA replication stress-induced transcriptome of Human Burkitt’s lymphoma identifies MBD1 as a novel suppressor of *BCL6* rearrangements in germinal center derived B-lymphomagenesis

**DOI:** 10.1101/2025.03.13.643172

**Authors:** Santosh Kumar Gothwal, Kyoko Oichai, Jacqueline H Barlow

## Abstract

BCL6 is a master transcriptional regulator of germinal center (GC) B cells. *BCL6* is frequently translocated at the major translocation cluster (MTC) within intron 1 of the *BCL6* locus, a hotspot commonly rearranged in diffuse large B cell lymphomas (DLBCLs). *BCL6* amplifications are associated with therapeutic resistance and poor survival outcomes in hematological and solid cancers. However the mechanisms suppressing genome instability at the *BCL6*-MTC preventing *BCL6* rearragements remain unclear. Here, transcriptome analysis and genome-wide mapping of histone H3 lysine 4 trimethylation (H3K4me3) in hydroxyurea (HU)-treated Raji cells (a Burkitt’s lymphoma model) revealed the induced expression of *MBD1*, encoding the DNA CpG methylation-binding protein. Functional studies using shRNA silencing and ectopic overexpression demonstrated that MBD1 suppresses *BCL6* transcription whose promoter harbours conserved CpG methylation sites, suggesting a DNA methylation-dependent regulation of *BCL6* trasncription by MBD1. Conversely, BCL6 repressed *MBD1* expression by binding to its promoter. *MBD1*-depleted Raji cells exhibited increased genomic instability at the *BCL6*-MTC upon HU treatment, heightened sensitivity to DNA replication inhibitors (HU, gemcitabine, and etoposide), and reduced tumorigenicity in xenograft mouse models. We propose that MBD1 prevents genomic instability at the *BCL6*-MTC to suppress DLBCL formation. Moreover, MBD1 promotes genomic stability and cell viability during DNA replication stress. MBD1 thus represents a potential therapeutic target for cancers exhibiting resistance to chemotherapies targeting DNA replication.

**Key Points:**

1. MBD1 suppresses *BCL6* expression under the DNA replication stress
2. MBD1 suppresses genomic instability at *BCL6* translocation hotspots
3. MBD1 depletion is associated with reduced tumorigenicity in mouse xenograft and sensitivity to DNA replication inhibitors.

## Introduction

Dark zones (DZs) and Light zones (LZs) are two main anatomical compartments (Phan, 2005 #36) within GCs [1]. DZs accommodate GC B cells undergoing activation induced cytosine deaminase (AID)-induced somatic hypermutation (SHM) [2], and LZs are proposed anatomical sites favoring AID induced class switch recombination (CSR) [3, 4]. GC B cells remain at high risk of genetic rearrangement in both DZs and LZs because dysregulated AID activity during SHM and CSR can initiate oncogenic chromosomal translocations including those fusing the Immunoglobulin locus (*IG*) with trans loci such as *MYC* and *BCL6* [5, 6]. GC B cells with these translocations develop into Burkitt’s lymphoma *(IG-MYC*) and DLBCLs (*IG-BCL6*) [5]. The hijacked regulation of differentiation pathways toward plasma and memory B cells in Germinal center derived B-lymphomas (GCDBL) can lead to Multiple Myeloma (MM) and B cell chronic lymphocytic leukemia (B-CLL), respectively [7]. Additionally, DLBCLs overexpressing *CXCR4*, which encodes G-protein coupled receptors (GPCRs), can develop into Waldenström macroglobulinemia, another DLBCL subtype marked by *BCL6* and *CXCR4* co-amplification [1, 8–11]. Thus, controlling genetic rearrangements in GC B cells, along with regulated differentiation and affinity maturation, is important for GCDBL elimination.

The BCL6 is a master transcriptional regulator of GC B cells belonging to the zinc finger and BTB (ZBTB) family and is comprised of an N-terminal BTB/POZ (Broad-complex, Tramtrack, Bric-a-brac/Poxvirus and zinc fingers) domain and *Krüppel*-type zinc fingers at the C terminus [12, 13]. The zinc finger domain of BCL6 is required for DNA binding to recruit the SMRT/mSIN3A/histone deacetylase for transcriptional suppression of genes in cell cycle arrest, DNA damage repair and p53 response [12, 13]. Thus, BCL6 drives the transcription program promoting GC B cell activation and survival by inhibiting the DNA damage response, apoptosis in GC B cells undergoing the SHM and CSR [14, 15].

*BCL6* overexpression is frequently observed in DLBCL [5, 16]. The breakpoint analysis of hematological malignancies suggests that 31% of DLBCLs harbor *BCL6* translocations on intron 1, a 10502 base pair sequence also known as major translocation cluster (MTC) [5, 16]. The start codon of *BCL6* is located on exon 3, thus translocations involving intron 1 encodes for full length BCL6 when juxtaposed with an active locus in trans. 52% of *BCL6* translocations involve the Immunoglobulin heavy chain (*IGH*) locus, 10% with immunoglobulin light chain (*IGL*), and 38% with non-*IG* loci [5, 16, 17], suggesting random nature of *BCL6*-translocation to sites other than *IG* locus. BCL6 also acts as a proto-oncogene in the pathogenesis of breakpoint cluster region (*BCLR*)*–*v-abl Abelson murine leukemia viral oncogene homolog 1(*ABL1*)-driven acute lymphoblastic leukemia (ALL). *BCL6* translocations have been also observed in glioblastoma and elevated BCL6 activity—along with its corepressor NcoR—is associated with AXL signaling and therapy resistance in glioblastoma [18]. BCL6 overexpression enables survival of Philadelphia chromosome-positive ALL (Ph^+^ ALL) cells upon BCLR–ABL1–kinase inhibition through repression of p53 expression [19]. BCL6 also drives therapy escape in non-small cell lung cancers, breast cancer, colorectal and gall bladder cancer [20, 21].

GC B cell differentiation is driven by dynamic changes in chromatin state including the DNA methylation and EZH2 driven histone H3K27me3 of the genes regulating the plasma B cell differentiation such as *IRF4* and *PRDM1* [22, 23]. The DNA CpG methylation binding protein 1 (MBD1) exhibits binding to methylated and unmethylated DNA via its CXXC domains ({Fujita, 2003 #78) and regulates gene expression by recruiting transcriptional suppressors such as SETDB2, HP-1α, SUV39H, histone-deacetylases-3 and PML-RARA {Fujita, 2003 #78;Clouaire, 2010 #79[24]. However, it is not yet clear if MBD1 regulates *BCL6* transcription through DNA methylation-dependent or -independent manner. During the plasma B cell differentiation, DNA hypomethylation of the promoters of *PRDM1, XBP1, IRF8, SPIB* genes assists plasma B cell differentiation [22, 25]. DNA methyltransferase activity was reported to be decreased in GC B cells than naïve B cells and *Dnmt1* hypomorphic mice had a defective GC reaction [22], suggesting a role for DNA methylation axis in GC B cell activation and differentiation.

Intron 1 of *BCL6* locus harbors four major CpG rich domains [16] [5], suggesting a role of DNA methylation in *BCL6* transcription. CTCF, an insulator which binds to unmethylated DNA, also binds on intron 1 and suppresses *BCL6* transcription, suggesting this transcriptional suppression is methylation-independent [26]. CpG methylation is catalyzed by the ten eleven translocation (TET) enzymes and is recognized by methylation binding proteins (MBDs), suggesting a complex regulation involving DNA demethylation, TET enzymes, MBD and CTCF binding in the regulation of *BCL6* transcription [4, 27–30]. The mechanistic link between DNA methylation-dependent suppression of *BCL6* transcription and whether MBD proteins are involved in *BCL6* transcription and remain unknown.

In the current study, we have characterized the roles of MBD1 in regulation of key genes involved in GC B cell fate determination utilizing a recently described approach for identification of new GC regulators. Functional characterization of MBD1 revealed its roles in suppression of *BCL6* under DNA replication stress. Moreover, *MBD1* knockdown resulted in increased DNA damage on the *BCL6*-MTC region while MBD1-depelted Raji cells exhibited higher sensitivity to DNA replication inhibitors. These place MBD1 as a novel regulator of BCL6 translocation and expression of *BCL6*. This role of MBD1 in GC B cells undergoing dynamic chromatin modification and differentiation could be essential for allowing the differentiation of GC B cells assuring the healthy immune response and suppression of the GCDBLs.

## Results

### Inverse correlation between *MBD1* and *BCL6* expression in GC B and cancers

We recently defined a novel approach using Raji cells to define the molecular mechanisms regulating the fate of GC B cells and GCDBLs [31]. Briefly, we exposed Raji cells to 4mM HU for 12 hours to induce the genotoxic-stress as experienced by GC B cells experience during rapid proliferation coupled with SHM and CSR [31]. We hypothesized that factors induced during this treatment could be important regulators on GC B cells. Based on this screening, we centered our investigation on *MBD1* due to its significant upregulation as one the the top 50 genes identified in our study (Figure 1A). We first confirmed whether *MBD1* expression is induced in cancer cells treated with DNA replication inhibitors and the DNA damage inducers Hydroxyurea (HU), Gemcitabine, and Etoposide (Supplementary Figures 1A, B). We performed short hairpin RNA (shRNA) mediated knockdown of *MBD1* in Raji cells (shMBD1-Raji) and in breast cancer cells (shMBD1-SUM149PT), and control cells with scrambled shRNA (shSCR-Raji and shSCR-SUM149PT) (Supplementary Figures 1A, B). We confimed that *MBD1* expression was significantly induced in scramble-knockdown cells, shSCR-Raji cells treated with HU, Gemcitabine or Etoposide (Supplementary Figures 1A). While we confirmed that shMBD1 cells reduced MBD1 expression by ∼75% compared to shSCR-Raji cells (Supplementary Figure 1A), shMBD1-Raji cells treated with these agents also exhibited an trend of induced MBD1 expression (Supplementary Figure 1A). Similarly, breast cancer origin cells, shSCR-SUM149PT also induced *MBD1* expression with HU-treatment (Supplementary Figure 2B). These results suggest that *MBD1* expression is induced in DNA replication stress-dependent manner. In addition, we observed histone H3K4me3 enrichment within -2 kilobase pairs of its TSS in HU-treated cells (Figure 1B), suggesting MBD1 expression is partially dependent on histone H3K4me3 under HU stress (Figure 1B).

**Figure 1:**
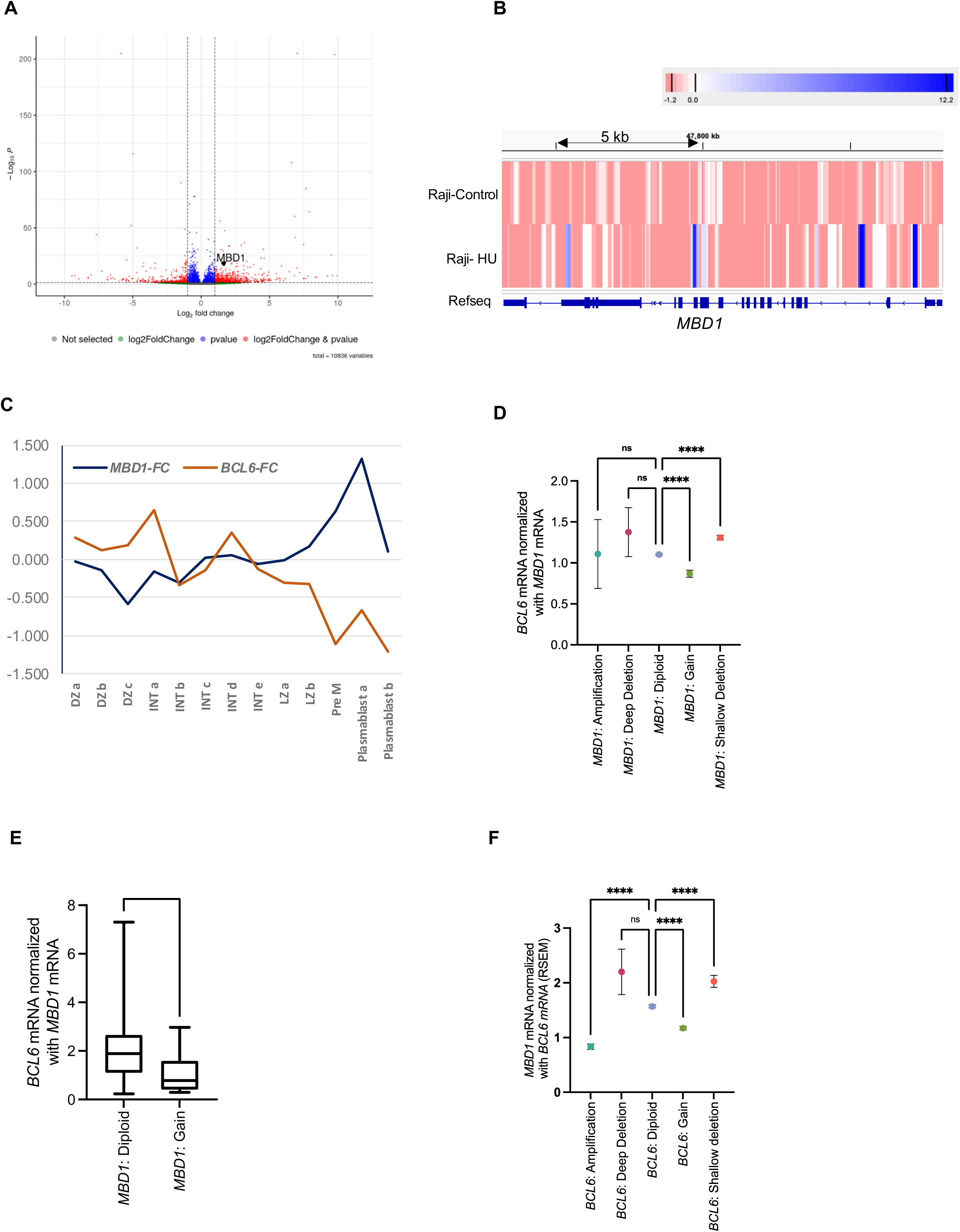
Identification of the DNA-replication stress-induced transcriptome in Raji cells. **(A)** Differentially expressed genes in control and HU-treated Raji cells (n=3) x axis indicated log2FC for mRNA fold change and y axis indicated -log10 of adjusted p values. data is from three independent samples **(B)** Analysis of histone H3K4me3 trimethylation ChIP-seq on the *MBD1* locus in vehicle-treated or HU-treated Raji cells **(C)** Real-time kinetics of *MBD1* and *BCL6* expression in activated germinal center B cells (AGCBs) from human tonsil **(D)** *BCL6-*diploid diploid tumors were classified into *MBD1*-amplification (n=6), *MBD1*-deep deletion (n=19), *MBD1*: diploid (n=4230), *MBD1*:gain (n=298) and *MBD1*-shallow deletion (n=1397) exhibiting mean *BCL6*-mRNA values of 1.110, 1.375,1.103, 0.8686 and 1.309 in each group respectively. p=<0.0001 *MBD1*:diploids vs *MBD1*:gain and *MBD1*-diploid vs MBD1-shallow deletion respectively. Dunnett’s multiple comparison test One way ANOVA. Data are presented as mean ± SEM. **(E)** Comparison of *BCL6* mRNA levels in the DLBCL samples (TCGA database) exhibiting either *MBD1*:diploid or *MBD1*:gain status. Normalization of *BCL6* mRNA in MBD1 altered groups was calculated by normalizing the RSEM values of *BCL6* mRNA by RSEM values of *MBD1* mRNA in each sample. p= 0.0.0307 for *MBD1*:diploid vs *MBD1*:gain. T-test, Mann-Whitney test **(F)** *MBD1*-diploid tumors were classified into *BCL6*-amplification, *BCL6*-deep deletion, *BCL6*-diploid, and *BCL6*-gain and *BCL6*-shallow deletion groups. Values on y-axis indicates normalized RSEM values of *MBD1* mRNA with the RSEM values of *BCL6* mRNA. The mean of normalized *MBD1* mRNA was 0.8332 in *BCL6*-amplification (n=182), 2.201 for *BCL6*-deep deletion (n=15), 1.57 in *BCL6*-diploid (n=4230), 1.171 in *BCL6*-gain (n=972), and 2.027 in *BCL6*-shallow deletion (n=390) group. *p< 0.0001* for *BCL6*-amplification vs *BCL6*-diploid, *p< 0.0001* for *BCL6*-diploid vs *BCL6*-gain. Dunnett’s multiple comparison test, One way ANOVA.

To examine the expression of *MBD1* in GC B cells, we analyzed the real-time transcriptome data from activated human tonsil GC B cells (AGCBs) (Figure 1C) [32]. We simultaneously plotted the log2 fold change (Log2FC) values of *MBD1* transcripts against *BCL6* transcripts across subpopulations, including DZa, DZb, DZc, and intermediate (INT) populations (INTa, INTb, INTc, INTd, INTe), as well as LZa, LZb, LZc, pre-memory (preM), and plasmablasts A and B (Figure 1C). These subpopulations were clustered based on genes specifically enriched in each compartment [32]. Log2FC values for *MBD1* were relatively low in DZa, DZb, INTa, INTb, INTc, INTd, INTe, LZa, and LZb, with values ranging from -0.34 to 0.163 (Figure 1C). However, log2FC values of *MBD1* were higher in pre-memory (0.625) and plasmablast a cells (1.320) (Figure 1C). In contrast, *BCL6* expression sharply declined in pre-memory and plasmablast stages (Figure 1C), suggesting a reciprocal relationship between *MBD1* and *BCL6* expression in GC B cells.

Given the reciprocal expression of *BCL6* and *MBD1* in human AGCBs, we checked the possibility of mututal suppression of MBD1 and BCL6. Among the tumor samples available in the cancer genome atlas (TCGA), we first categorized *BCL6* and *MBD1* altered tumors based on their genetic status and quantified their mRNA level in each group (Supplementary Figure 1B, C). We found that *MBD1* mRNA levels positively correlate with genetic alteration (Supplementary Figure 1C). We also observed a similar relationhip of *BCL6* mRNA expression with its copy number aleration in tumors (Supplementary Figure 1D).

We then examined *MBD1*-mutated tumors for alterations in *BCL6* expression. Tumors with *MBD1*-shallow deletions exhibited higher *BCL6* expression than *MBD1*-diploid tumors (Figure 1D). On the other hand, *BCL6* mRNA levels were significantly lower in *MBD1-gain* tumors (Figure 1D). Similarly, *MBD1*-amplified tumors exhibited reduced trend of *BCL6* mRNA expression while tumors with *MBD1*-deep-deletion status exhibited higher *BCL6* mRNA levels than *MBD1*-diploid tumors, although this trend was not significant in these groups (Figure 1D). Further, DLBCL samples with *MBD1-gain* status exhibited lower *BCL6* expression than *MBD1-* diploid tumors (Figure 1E). Taken together, these results indicate an inverse relationship between *MBD1* and *BCL6* expression. To determine if *BCL6* levels also inversely correlate with *MBD1* mRNA levels, we sorted tumors with *MBD1*-diploid samples and classified them as *BCL6*-diploid, *BCL6*-deep-deletion, *BCL6*-shallow-deletion, *BCL6*-gain and *BCL6*-amplification (Figure 1F). We found that *MBD1* mRNA was significantly lower in *BCL6*-gain and *BCL6*-amplified cancers, while *MBD1*-mRNA was higher in *BCL6*-deleted samples (Figure 1F). These results indiciate that *BCL6* levels are inversely corealted with the *MBD1* mRNA levels, sugesting the mutual suppresion of BCL6 and MBD1 by each others.

### MBD1 suppresses *BCL6* expression during DNA replication stress

Given the abnormal expression of *BCL6* in breast cancer cells, we investigated whether *MBD1* knockdown also impacts *BCL6* levels in breast cancer cell lines MCF7, MDA-MB-231, SUM-149PT, and MDA-MB-436 (Figure 2A). Notably, *BCL6* expression was induced in shMBD1-MCF7 and shMBD1-MDA-MB-231 cells compared to control cells (Figure 2A). These results confirm that MBD1 can suppress *BCL6* expression (Figure 2A). However, *MBD1* depletion in SUM149PT and MDA-MB-436 cells did not induce the *BCL6* suppression, suggesting additional mechanisms may regulate *BCL6* transcription in these cells. Overall, these results confirm that MBD1 loss is associated with the induced *BCL6* expression not only in B-lymphoma cells but also in breast cancer.

**Figure 2:**
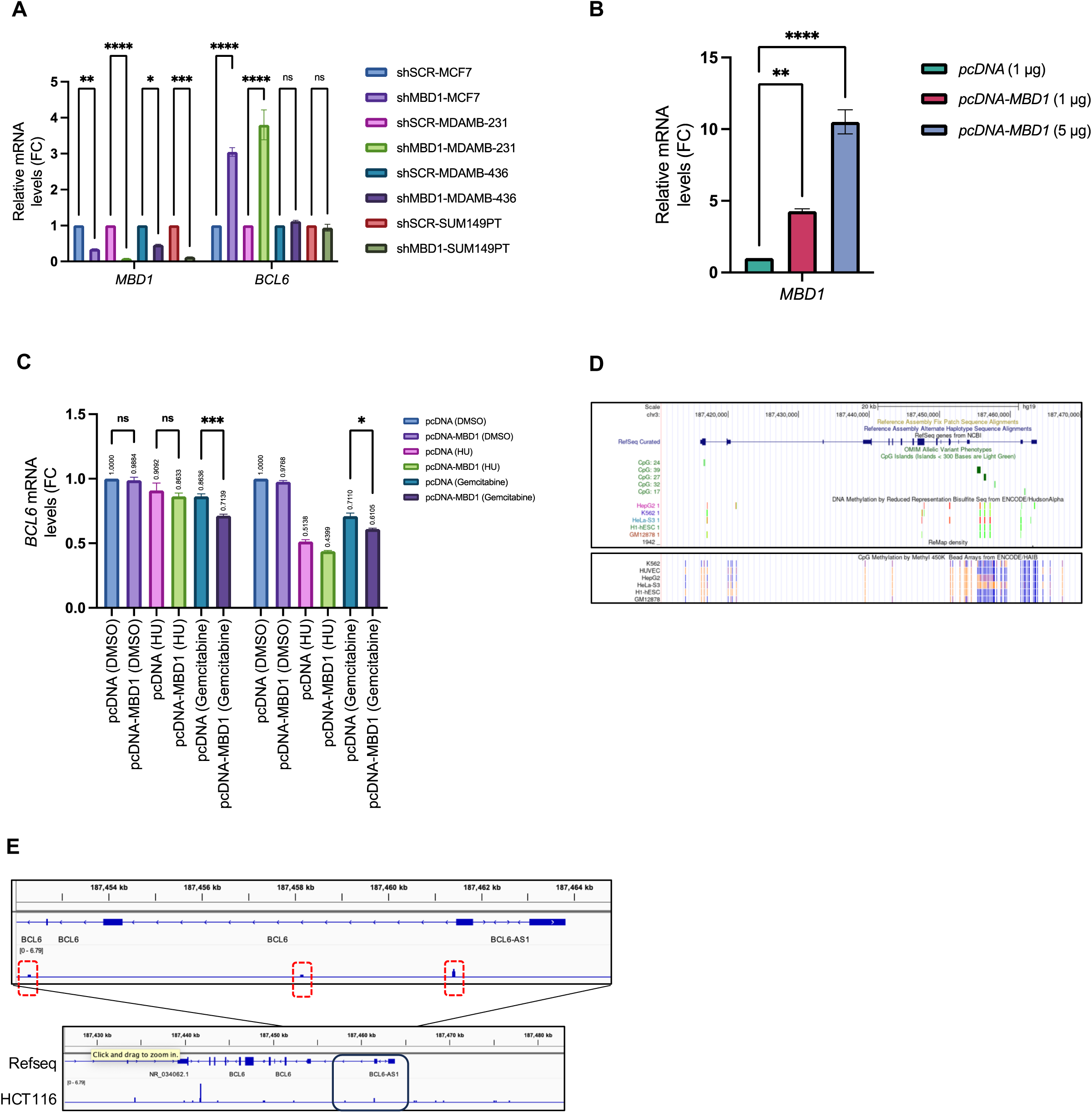
MBD1 suppresses *BCL6* expression under DNA replication stress. **(A)** shMBD1-MCF7, and shMBD1-MDA-MB-231 cells exhibit higher levels of *BCL6* expression than shSCR-MCF7 and shSCR-MDA-MB-231 cells. For *BCL6* group: *p<0.0001* for shSCR-MCF7 vs shMBD1-MCF7 cells., *p<0.0001* for shSCR-MDA-MB-231 vs shMBD1-MDA-MB-231 cells, p= 0.9927 for shSCR-MDAMB436 vs shMBD1-MDAMB436, p>0.9999 for shSCR-SUM149PT vs shMBD1-SUM149PT. For *MBD1*: p=0.0051 for shSCR-MCF7 vs shMBD1-MCF7, p<0.0001 for shSCR-MDAMB231 vs shSCR-MDAMB231, p=0.0339 for shSCR-MDAMB436 vs shMBD1-MDAMB436, p=0.0001 for shSCR-SUM149PT vs shMBD1-SUM149PT. Two-way ANOVA. Tukey’s multiple comparison test **(B)** *pcDNA* (1 μg) or *pcDNA-MBD1* (1 μg and 5 μg) were transfected into 0.5 million Raji cells seeded in 12-well plates. RNA was extracted 24 hours post-transfection, and cDNA preparation and qPCR were conducted for *MBD1.* p=0.0070 for *pcDNA* (1 μg) vs *pcDNA-MBD1* (1 μg) and p<0.0001 for *pcDNA* (1 μg) vs *pcDNA-MBD1* (5 μg). Sidak’s multiple comparison test (ordinary one-way Anova). Data are presented as mean ± SEM from three independent experiments **(C)** *MBD1* overexpression was performed in Raji cells and cells were treated with HU and gemcitabine for 24 and 48 hours followed by estimation of *BCL6* mRNA levels by qRT-PCR. For 24 hour-treated samples: p=0.9990 for *pcDNA* (DMSO) vs *pcDNA-MBD1* (DMSO), p=0.6897 for *pcDNA* (HU) vs *pcDNA-MBD1* (HU), and p=0.0009 for *pcDNA* (Gemcitabine) vs *pcDNA-MBD1* (Gemcitabine). For 48 hour-treated samples: p=0.9748 for *pcDNA* (DMSO) vs *pcDNA-MBD1* (DMSO), p=0.2108 for *pcDNA* (HU) vs *pcDNA-MBD1* (HU), and p=0.0394 for pcDNA (Gemcitabine) vs *pcDNA-MBD1* (Gemcitabine). Two-way ANOVA. Tukey’s multiple comparison test **(D)** *BCL6* promoter harbours CpG Methylation sites. Encode datasets of CpG methylation on *BCL6* gene among several cells lines is shown using the UCSD datasets **(E)** MBD1 binds to the *BCL6* promoters in HCT116 cells. Region highlighted in the above panel indicates the CpG methylation-rich regions exhibiting MBD1 peaks in HCT116. The lower panel shows *BCL6* genetic locus with MBD1 peaks through the gene body. MDB1 binding on *BCL6* locus was examined using the dataset #DRX021118 from the ChIP-atlas. (https://chip-atlas.org/view?id=DRX021118).

To further confirm that MBD1 suppresses *BCL6* expression in B cells, we transiently transfected Raji cells with *pcDNA-MBD1* (Figure 2B, C). Compared to *pcDNA3.1* (empty vector: EV) transfected cells, *pcDNA-MBD1* cells induced *MBD1* expression in Raji cells (Figure 2B). While *pcDNA-MBD1*-transfected cells did not have significantly reduced *BCL6* mRNA, treatment with gemcitabine caused a significant reduction in *BCL6* levels in *pcDNA-MBD1* transfected cells (Figure 2C). In addition, 24 and 48 hours of gemcitabine treatment yielded significant reduction in *BCL6* mRNA since EV-transfected Raji cells exhibited 86% and 71% of *BCL6* levels after 24 and 48 hours of gemcitabine treatment, respectively, while *pcDNA-MBD1*-transfected Raji cells showed 71% and 61% of *BCL6* levels (Figure 2C). Exposure to HU induced a similar trend: after 24 hours of HU treatment, *BCL6* levels were 90% in EV-transfected Raji cells and 86% in *pcDNA-MBD1*-transfected cells (Figure 2C). Though not significant, this trend persisted after 48 hours of HU treatment, with *pcDNA-MBD1* showing 43% *BCL6* mRNA levels compared to 51% in EV-transfected cells (Figure 2C). These results further support a role for MBD1 in suppressing *BCL6* transcription during replication stress. This is distinct from breast cancer cells, where exogenous DNA replication stress was not required for MBD1-mediated suppression of *BCL6* expression (Figure 2A).

MBD1 recognizes DNA methylation on CpG sequences[24, 33, 34], we investigated whether MBD1 might bind to methylated DNA sequences on the *BCL6* locus and suppress *BCL6* transcription. Using the ENCODE database, we identified four dispersed CpG methylation sites within intron 1 of the *BCL6* locus, where the *BCL6*-MTC resides (Figure 2D). In addition, by analyzing ChIP-atlas data and MBD1 ChIP-seq results from HCT116 cells, we confirmed MBD1 binding along the *BCL6* gene body including intron 1 (Figure 2E). These results suggest that MBD1 binds to the *BCL6* locus and the repression of *BCL6* transcription could be influenced by DNA methylation and DNA replication stress.

### MBD1 suppresses DNA break formation on the *BCL6*-MTC in Raji cells-

Mutations increasing *MBD1* expression negatively correlate with *BCL6* transcription (Figure 1D), however the relationship between *MBD1* and *BCL6* mutations in tumors is unclear. To determine if *MBD1* and *BCL6* mutations co-occur, we analyzed their co-occurrence in 27 different tumor types using TCGA (Figure 3A). We found that *MBD1* deep-deletion or *MBD1*-shallow deletion samples (commonly referred as *MBD1*-deleted hereafter) exhibited a higher percentage of *BCL6* alterations, suggesting that genetic alterations in *MBD1* and *BCL6* are positively associated (Figure 3A). Further, DLBCL tumors with *MBD1*-deleted status exhibited a significantly higher frequency of *BCL6* alterations, suggesting a strong correlation between *MBD1* and *BCL6*-mutated B lymphomas (Figure 3B). Based on these results, we hypothesized that MBD1 may also influence *BCL6* translocation by suppressing DNA breaks on *BCL6*-MTC preventing its rearrangement. To investigate this hypothesis, we analyzed γH2AX signal by chromatin immunoprecipitation (ChIP) on *BCL6*-MTC after HU-induced DNA replication stress in shSCR-Raji and shSCR-MBD1 cells (Figure 3C). We analyzed γH2AX abundance at four primer locations G, J, M and P on *BCL6* intron 1, which spans 10.5 kb (Figure 3C) [26]. The level of γH2AX signal was higher in HU-treated shMBD1-Raji cells than HU-treated shSCR-Raji cells (Figure 3D), suggesting that MBD1 prevents HU-induced DNA damage at *BCL6*-MTC (Figure 3D). This role of MBD1 in suppressing DNA damage at *BCL6*-MTC suggests that it could suppress *BCL6* rearrangements in GC B cells and prevent DLBCL formation. Since CTCF binds to intron 1 of *BCL6*-MTC and suppresses *BCL6* transcription in MM cells [26], we hypothesized that MBD1 may prevent *BCL6* transcription by collaborating with CTCF. We compared CTCF ChIP signal in HU-treated shMBD1-Raji and shSCR-Raji cells on the *BCL6*-MTC (Figure 3E). Compared to HU-treated shSCR-Raji cells, HU-treated shMBD1-Raji cells exhibited decreased CTCF occupancy at *BCL6*-MTC compared to HU-treated shSCR-Raji cells (Figure 3E). This result suggests that MBD1 promotes CTCF recruitment to *BCL6*-MTC during replication stress (Figure 3E).

**Figure 3:**
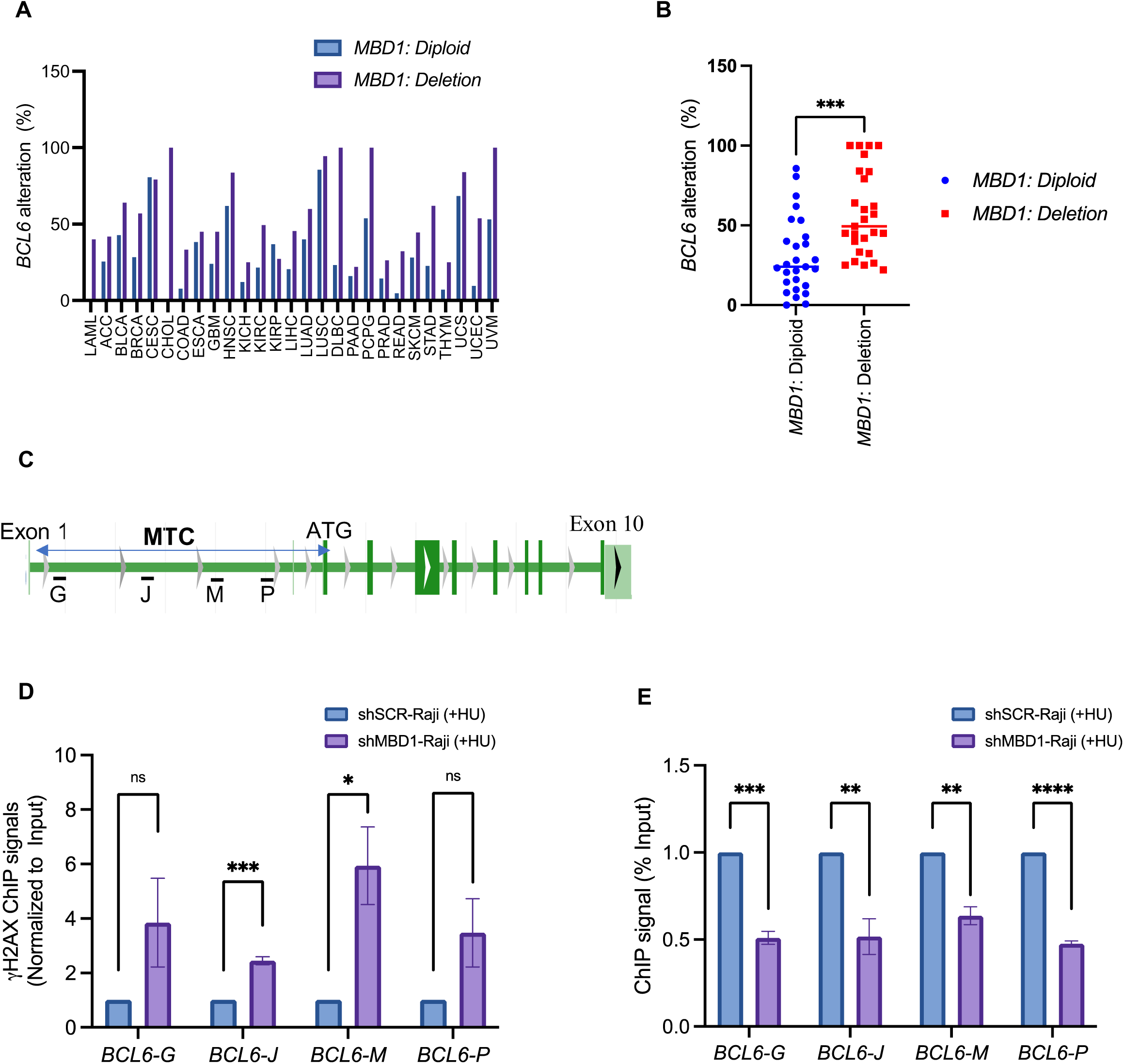
Increased genomic instability at *BCL6*-MTC in *MBD1*-depleted cells. **(A)** TCGA dataset of 27 tumor types exhibiting the *MBD1*-diploid of *MBD1*-deletion status. Deletion and shallow deletion samples are counted as deletion samples. Percentage of *BCL6* alteration is compared in each tumor type between *MBD1*:diploid or *MBD1*:deletion status. *BCL6* alteration are the samples exhibiting with *BCL6*:deep deletion, *BCL6*:shallow deletion, *BCL6*:gain or *BCL6*:amplification. Tumor samples exhibiting *BCL6*:diploid status are not included as altered *BCL6* status **(B)** The percentage of *BCL6* alterations in DLBCL samples from TCGA database was quantified in *MBD1*-diploid and *MBD1*-deleted tumors and presented in the graph. *MBD1*-deleted tumors include samples with *MBD1* deep and shallow-deletions. *BCL6* alterations encompass tumor samples with *BCL6* deep-deletion, shallow-deletion, gain, and amplification, while *BCL6* diploid samples were not considered altered **(C)** Schematic representation of the major translocation cluster of the human *BCL6* gene indicating exons and introns. Primer pair G, J, M, P location analyzed for ChIP-qPCR is indicated **(D)** γH2AX ChIP analysis in shSCR-Raji cells and shMBD1-Raji cells treated with 10 mM HU. qPCR analysis was performed using four primer pairs, G, J, M, and P, targeting intron 1 of the *BCL6* locus. Primer pair locations are not to scale. For primer pair G, *p = 0.1563* for shSCR-Raji (10 mM HU) vs. shMBD1-Raji (10 mM HU); for J, *p = 0.0006* for shSCR-Raji (10 mM HU) vs. shMBD1-Raji (10 mM HU); for M, *p = 0.0259* for shSCR-Raji (10 mM HU) vs. shMBD1-Raji (10 mM HU); for P, p = *0.1209* for shSCR-Raji (10 mM HU) vs. shMBD1-Raji (10 mM HU). Unpaired t-test. Data are presented as mean ± SEM (n=3) **(E)** CTCF ChIP analysis in vehicle- or HU-treated shSCR-Raji cells and HU-treated Raji cells. For primer pair G, p = 0.0002 for shSCR-Raji (10 mM HU) vs. shMBD1-Raji (10 mM HU); for J, *p = 0.0093* for shSCR-Raji (10 mM HU) vs. shMBD1-Raji (10 mM HU); for M, *p = 0.0022* for shSCR-Raji (10 mM HU) vs. shMBD1-Raji (10 mM HU); for P, *p <0.0001* for shSCR-Raji (10 mM HU) vs. shMBD1-Raji (10 mM HU) Unpaired t-test. Data are presented as mean ± SEM (n=3).

### BCL6 negatively regulates *MBD1* transcription

BCL6 suppresses plasma B cell differentiation by inhibiting expression of *IRF4* and *PRDM1* [35]. Given the inverse correlation of *BCL6* and *MBD1* expression in AGCBs (Figure 1C, D, F), we hypothesized that BCL6 may suppress *MBD1* expression in GC B cells undergoing plasma B cell differentiation. To test the effect of BCL6 inhibition on *MBD1* expression, we treated Raji cells with FX1, a potent BCL6 inhibitor which binds to the BCL6 lateral groove and inhibits the formation of the BCL6 repression complex [36]. We found that FX1 treatment significantly induced *MBD1* expression in Raji cells (Figure 4A), suggesting that inhibition of BCL6 binding to its DNA target sequences suppresses *MBD1* transcription (Figure 4A). These results suggest that MBD1 and BCL6 can regulate each other’s expression in GB B cells and is suggestive of mutual negative feedback. In addition, shBCL6-Raji cells exhibited a trend of increased *MBD1* expression compared to *shSCR-Raji* cells, however this difference was not significant (Figure 4B). We confirmed that *BCL6* knockdown was significantly achieved in shBCL6-Raji cells (Figure 4B), and *IRF4* levels were enhanced in shBCL6-Raji cells confirming the previous reports on BCL6 of the *IRF4* [37].

**Figure 4:**
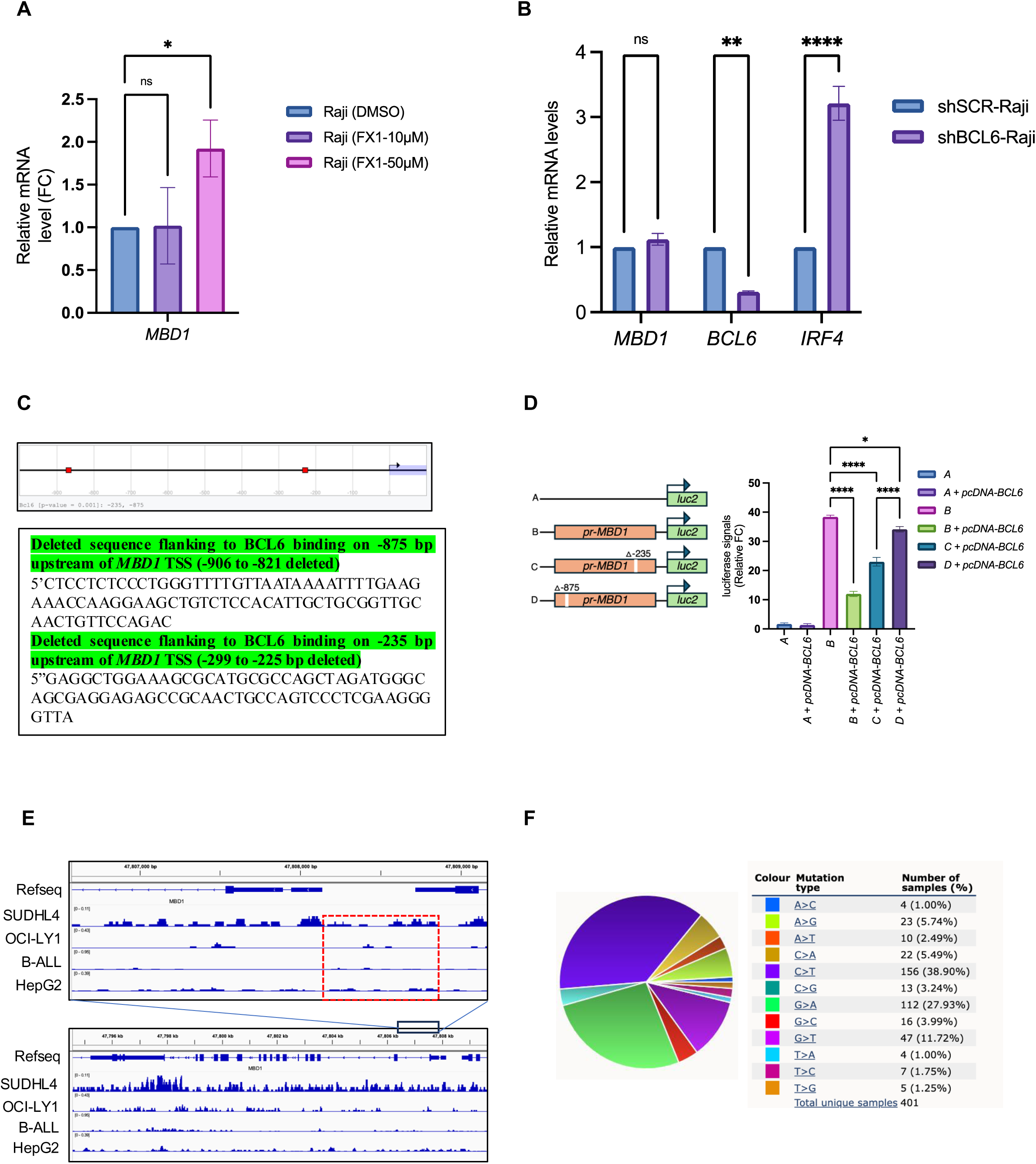
BCL6 and AID inactivate *MBD1* at transcriptional and genetic levels. **(A)** Treatment of Raji cells with the BCL6 inhibitor, FX1 (10 μM and 50 μM), induces MBD1 transcript levels in a dose-dependent manner. Data are presented as mean ± SEM from three independent experiments. Dunnett’s multiple comparison test: *p = 0.0223* for Raji (DMSO) vs. Raji (FX1-50 μM) treatment groups **(B)** qPCR analysis of *BCL6*, *MBD1*, and *IRF4* transcript levels in shSCR-Raji and shBCL6-Raji cells. Sidak’s multiple comparison test: two-way ANNOVA, For *MBD1*, *p =0,8412* for shSCR-Raji vs shBCL6-Raji groups; For *BCL6,* p=0.0030 for shSCR-Raji vs. shBCL6-Raji. For *IRF4* group, p <0.0001 for shSCR-Raji vs. shBCL6-Raji groups **(C)** In silico prediction of the BCL6 binding motif at the *MBD1* promoter using JASPAR CORE 2018 Vertebrates, available in the Eukaryotic Promoter Database (EPD). The sequences of two BCL6 binding motif located on -235 and -875 base pair upstream of *MBD1* TSS are indicated. The given sequence represents the regions deleted in the *pGL4-pr-MBD1* plasmids **(D)** Schematic representation of the *MBD1* promoter cloned upstream of the luciferase gene in *pGL4-luc2* and the BCL6 binding motif. Location of BCL6 binding motif and deletion are shown. HEK293T cells were transfected with *pGL4-luc2*, *pGL4-luc2 + pcDNA-BCL6*, *pGL4-luc2-pr-MBD1*, *pGL4-pr-MBD1* + *pcDNA-BCL6*, *pGL4-luc2-pr-MBD1Δ235* + *pcDNA-BCL6*, and *pGL4-luc2-pr-MBD1Δ900* + *pcDNA-BCL6* plasmids and examined for luciferase activity after 24 hours. Data are presented as mean ± SEM from three independent experiments. Tukey’s multiple comparison test: *p < 0.0001* for *pGL4-luc2* vs. *pGL4-luc2-pr-MBD1*; *p < 0.0001* for *pGL4-luc2-pr-MBD1* vs. *pGL4-luc2-pr-MBD1* + *pcDNA-BCL6* groups; *p < 0.0001* for *pGL4-luc2-pr-MBD1* vs. *pGL4-luc2-pr-MBD1Δ235* + *pcDNA-BCL6*; *p = 0.0416* for *pGL4-luc2-pr-MBD1* vs. *pGL4-luc2-pr-MBD1Δ900* + *pcDNA-BCL6* **(E)** BCL6 binds to *MBD1* locus in SUDHL4, OCI-LY1 (DLBCL), B-ALL and HepG2 cells cells. The data are analyzed from ChIP-atlas database. Datasets: SUDHL4:SRX4609168; OCI-LY1:SRX689470, B-ALL:SRX18259603, HepG2:2636278 (https://chip-atlas.dbcls.jp/data/hg38/target/BCL6.10.html) **(F)** The COSMIC mutation landscape of the *MBD1* gene, showing the frequencies of A>C, A>G, A>T, C>A, C>T, C>G, G>A, G>C, G>T, T>A, T>C, and T>G mutations.

To explore how BCL6 suppresses *MBD1* transcription, we performed *in-silico* binding analysis of BCL6 on the *MBD1* promoter (Figure 4C). We observed two BCL6 binding motifs on the *MBD1* promoter present at -235 and -900 bp upstream of *MBD1*-TSS (Figure 4C, D). To determine if BCL6 binding occurs at these sites, we cloned 1 kb of the *MBD1* promoter region, and inserted into the luciferase reporter *pGL4* vector then performed site-directed mutagenesis on DNA sequences flanking these two binding sites (Figure 4D). 293T cells transfected with *pGL4-pr-MBD1* induced luciferase signal nearly 38-fold compared to 293T cells transfected with empty *pGL4* (Figure 4D). Interestingly, co-transfection of *pGL4-pr-MBD1* and *pcDNA-BCL6* reduced the luciferase signal to around 15%, suggesting that BCL6 suppresses *MBD1* expression by binding its promoter (Figure 4D). Deletion of either BCL6 binding motif significantly increased luciferase signal, suggesting that BCL6 binding to the predicted binding motifs negatively regulate transcription (Figure 4D). Of note, *pcDNA-BCL6* expression had a smaller impact on luciferase signal from the *pGL4-pr-MBD1 Δ235-BCL6* or *pGL4-pr-MBD1 Δ900-BCL6* reporters than the wild-type promoter sequence (Figure 4D). Particularly, deletion of the site located -900 upstream of *MBD1-TSS* (Figure 4D) had a greater rescue from BCL6-mediated suppression, suggesting that

BCL6 binding to this motif plays a crucial role in regulating MBD1 expression (Figure 4D). We next examined whether BCL6 binding is present on the endogenous *MBD1* promoter. Using BCL6 ChIP-seq datasets from ChIP-atlas, we confirmed BCL6 binding on *MBD1* promoter in multiple cell lines including SUDHL4 (DLBCL), OCI-LY1 (DLBCL), OCI-LY3 (DLBCL) and in B-cell acute lymphoblastic leukemia (B-ALL) cell lines (Figure 4E). These results suggest that BCL6 binds to the *MBD1* promoter and suppresses its transcription (Figure 4D, E).

To investigate whether BCL6 regulation is associated with *Mbd1* regulation in mouse GC B cells, we examined the DNA methylation status of the *Mbd1* promoter in wild-type and *Sca1-Bcl6^Δ^* mice (Supplementary Figure 2A[38]. We noted that the *Mbd1* promoter in activated GC B cells of wild type mice showed two distinct DNA methylation peaks, and one peak was lost in *Sca1-Bcl6^Δ^* mice (Supplementary Figure 2A). These results suggest that BCL6 suppression of *Mbd1* could be associated with DNA methylation on the *Mbd1* promoter in mouse GC B cells. To determine if methylation impacts *MBD1* expression, we analyzed human B lymphomas treated with 5- Azacytidine (AZA), a DNA methyltransferase inhibitor (Supplementary Figure 2B), and observed that AZA treatment increased transcripts per million (TPM) counts of *MBD1* in OCI-LY1, SUDHL2, and OCI-LY19 cells. Together these results suggest that MBD1 expression in GC B cells is regulated by BCL6 and DNA methylation (Supplementary Figure 2B).

GC B cells are regulated by BCL6 and AID, both of which also act as oncoproteins. Since BCL6 suppresses *MBD1* (Figure 4A, D), it is possible that MBD1 may function as a tumor suppressor in GC B cells. AID is also involved in B-lymphomagenesis due to its mutagenic roles in GC B cells [10, 39], therefore we explored if AID is associated with *MBD1* mutagenesis. We looked for AID signatures in *MBD1*-mutant cancers in the Catalogue of somatic mutation in cancer cells (COSMIC) (Figure 4F). Interestingly, AID mutation signatures (C>T), which are initiated by cysteine deamination to uracil in non-replicating B cells, were frequent (38.9%) along the *MBD1* gene in *MBD1* mutant tumors (Figure 4F). Furthermore, C>G and C>A signatures, which are not as strongly associated with AID activity and could arise due to translesion synthesis and mismatch repair, were 3.4 % and 5.5%, respectively (Figure 4F). Among non-AID signature mutations in the *MBD1* gene, G>A were the most frequent (27.9%) among all samples (Figure 4F). These results suggest that *MBD1* is mutated by AID and underscore MBD1 as a pivotal target for both BCL6 and AID in promoting B lymphomagenesis.

### Increased sensitivity of *MBD1*-depleted cells to chemotherapeutics targeting DNA replication and reduced tumorigenicity in mouse xenograft model

The increased levels of γH2AX signal at the *BCL6*-MTC in HU-treated shMBD1 cells indicates increased susceptibility of *MBD1*-depleted cells to chemotherapeutic agents inducing replication stress (Figure 3D). To determine if shMBD1-Raji cells exhibit higher susceptibility to other chemotherapeutic agents, we treated shSCR-Raji and shMBD1-Raji cells with gemcitabine or etoposide (Figure 5A). Control ShSCR-Raji cells showed modest sensitivity to HU and Gemcitabine, and greater sensitivity to etoposide (Figure 5A). However, depletion of *MBD1* led to significantly greater sensitivity to all drugs tested: the viability in shMBD1-Raji cells was further reduced to 60%, 81.5% and 40.5% compared to shSCR-Raji cells treated with HU, gemcitabine or etoposide (Figure 5A). These results suggest that *MBD1*-deficient cells are more sensitive to genotoxic agents than shSCR-Raji and indicate a role of MBD1 in promoting cell viability in response to DNA replication stress/genotoxic stress.

**Figure 5:**
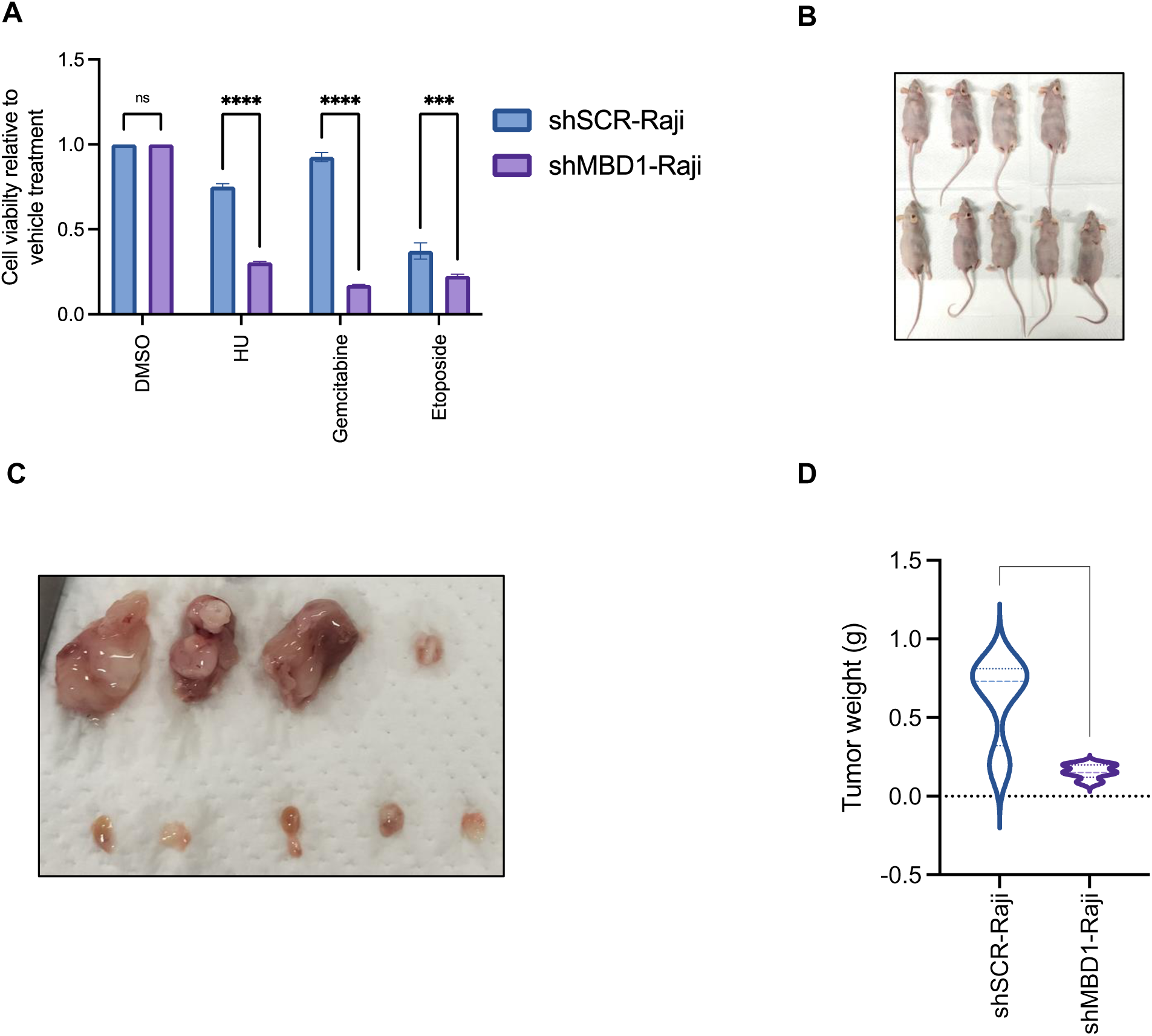
Reduced tumorigenicity of MBD1 depleted Raji cells in mouse xenograft model. **(A)** Cell viability of shSCR-Raji and shMBD1-Raji cells treated with DMSO, HU (10 mM), Gemcitabine (20 mM), and Etoposide (50 μM) for 24 hours. Sidak’s multiple comparison test: p < 0.001 for shSCR-Raji (HU) vs. shMBD1-Raji (HU); p < 0.0001 for shSCR-Raji (Gemcitabine) vs. shMBD1-Raji (Gemcitabine); p = 0.0005 for shSCR-Raji (Etoposide) vs. shMBD1-Raji (Etoposide) **(B)** Tumor xenografts were established by subcutaneously injecting 1 million shSCR-Raji or shMBD1-Raji cells mixed with 200 μl of Matrigel above the right front limb of nude mice to reach the subcutaneous pocket. Mice were monitored for tumor growth over a period of 55 days. The sample size consisted of n=4 for the shSCR-Raji group (Upper panel) and n=5 for the shMBD1-Raji group (lower panel) **(C)** Tumor xenografts from shSCR-Raji and shMBD1-Raji cells injected into nude mice (n=4 for shSCR-Raji; n=5 for shMBD1-Raji) **(D)** Violin plot showing the weight range of tumors in shSCR-Raji and shMBD1-Raji groups. Unpaired t-test: *p = 0.0086*.

Finally, to test whether tumorigenicity is altered in *MBD1-*silenced cells, we performed xenograft experiments by injecting shSCR-Raji and shMBD1-Raji cells into nude mice (Figure 5B-D). Mice injected with shSCR-Raji cells exhibited mean tumor weight of 0.62 gm (Figure 5D). In contrast, the mean weight of shMBD1-Raji tumors was 0.15 gm (Figure 5D). This reduction in tumor weight of shMBD1-Raji xenograft was significantly lower than those in shSCR-Raji cells (Figure 5C, D). This result suggests that MBD1 is required for the tumorgenicity of Raji cells *in vivo* (Figure 5E, C, D). It is possible that accumulative genotoxic stress may surpass the threshold cells can tolerate in shMBD1-Raji cells therefore perturbing the cellular proliferation, tumorgenicity and tumor growth (Figure 5B,C, D).

### Correlation of *MBD1* and *IRF4* expression in B-Lymphoma

Human AGCBs induce *MBD1* expression in GC B cells undergoing differentiation into plasma and memory B cells while *BCL6* expression was reduced at similar stages, suggesting that the upregulation of MBD1 in differentiating GC B cells is correlated with *BCL6* downregulation (Figure 1C). We hypothesized that *MBD1* could regulate the expression of genes in GC B cells undergoing terminal differentiation. To define the relationship between *MBD1* mutation status and *IRF4* expression, we explored their correlation in samples from TCGA datasets and the Gene Expression Profiling Interactive Analysis (GEPIA) (Supplementary Figure 3A, B). *IRF4* expression was highest in DLBCL samples exhibiting *MDB1*:gain status, suggesting a positive correlation between *MBD1* and *IRF4* expression (Supplementary Figure 3A). Moreover, DLBCL tumors from GEPIA also exhibited a positive correlation in *MBD1* and *IRF4* (Supplementary Figure 3B). Since *IRF4* expression is induced in plasma B cells, we examined whether mouse plasma B cells exhibit higher *Mbd1* expression than activated GC B cells (Supplementary Figure 3C). Interestingly, *Mbd1* expression was increased in mouse plasma B cells, when IRF4 is high (Supplementary Figure 3C) [40], suggesting that *Mbd1* induction in plasma B cells is coordinated with *Irf4* expression. We also measured the effect of *MBD1* knockdown on the expression of two key genes determining GC B cell fate, *IRF4* and *PRDM1* (Supplementary Figure 3D). *MBD1* depletion in Raji cells resulted in increased expression of *IRF4* and *PRDM1*, indicating a negative regulatory role of MBD1 in their transcription (Supplementary Figure 3D). Conversely, overexpression of MBD1 led to a reduction in *IRF4* expression (Supplementary Figure 3E), suggesting that MBD1 plays a critical role in *IRF4* regulation.

Finally, we analyzed if *MBD1* depletion affects the expression of key factors regulating GC B cell activation and survival including essential homing receptors *(CXCR4*, *CXCR5*, *CCR7*, *CXCR4*, *CXCR5*, *S1PR2*, *P2RY8)*, genes encoding the components of migration machinery (*RAC1*, *RHOA)*, BCL6 targets, (*IRF7*, *IFNGR1*, *STAT1),* or genes encoding factors regulating BCL6 turnover, (*FBXO11*, *XBP1)* (Supplementary Figure 3F). Among the GC B cell receptors tested, shMBD1-Raji cells exhibited increased expression of *CCR7*, *P2RY8* and *STAT1* but reduced expression of *RAC1* compared to shSCR-Raji cells (Supplementary Figure 3F). The expression of BCL6 targets or regulators *S1PR2*, *RHOA*, *FBXO11*, *XBP1*, *IRF7* and *IGNGR1* was not altered upon in the shMBD1-Raji cells (Supplementary Figure 3F). Overall, these observations suggest that MBD1 can affect the position of B cells within the GC compartment by regulating the expression levels of *CCR7*, *CXCR5*, and *CXCR4* in cells belonging to T-B border (T-B), light zone (LZ), and dark zone (DZ), respectively. MBD1’s regulation of *RAC1* and *P2RY8* may also affect the dynamic migration of GC B cells, given a role of RAC1 and P2RY8 in GC B cell LZ-migration [41, 42].

## Discussion

### Mechanism of *MBD1* induction in GC B cells undergoing Plasma B Cell Differentiation

The increased *MBD1* expression we observed in HU-treated Raji cells correlates with elevated *MBD1* levels in human AGCBs classified as pre-memory and plasma B cells [32] (Figure 1C). Similarly, mouse GC B cells undergoing PC differentiation also exhibit increased *Mbd1* levels (Supplementary Figure 3C). This evidence highlights MBD1’s role in GC B cell differentiation in both humans and mice (Figures 1C, supplementary Figure 3C). The transition from GC B cells to plasma cells necessitates dynamic DNA hypomethylation in genes involved GC B differentiation [25, 43], suggesting that MBD1 expression could be induced by hypomethylation of its promoter. This is supported by our observations of DNA methylation loss in the *Mbd1* promoter in *Sca1-BCL6^Δ^* mice (Supplementary Figure 2A), and the induction of *MBD1* expression in the presence of the methylation inhibitor AZA in DLBCLs (Supplementary Figure 2B) [38]. MBD1 induction during plasma B cell differentiation could also be related to *BCL6* downregulation since we provide evidence that *BCL6* suppresses *MBD1* expression by binding to its promoter (Figures 4A, D). The increased *MBD1* levels in plasma B cells might also result from reduced EZH2-dependent silencing of the *MBD1* promoter, as this suppresion is removed in GC B cell undergoing differentiation [23]. In contrast, induction of *MBD1* in cancer cells undergoing chemotherapy with DNA replication inbibitors suggests a programmed activation of *MBD1* transcription which potentially regulates cell survival (Supplementary Figure 1A, B, Figure 5A-D). The detailed mechanims regulating MBD1-mediated cell survival should be explored in future studies to rationally design effective combination therapy for *MBD1*-inactivated cancers. *MBD1* alteration may also be an effective biomarker for tumor exhibiting higher *BCL6* and *IRF4* expression which also correlates with increased chemotherapeutic resistance [19, 44].

### *BCL6* translocations and reduced tumorigenicity in *MBD1*-depleted cancer cells

The *BCL6*-MTC contains highly conserved CpG methylated sites on it intron 1 (Figure 2D) [26], therefore MBD1 may bind to the *BCL6*-MTC in a DNA methylation-dependent manner (Figure 2D, E). MBD1 binding can help recruit histone deacetylases like SUV39H for transcriptional repression at the *BCL6* locus [5, 24, 33, 45]. MBD1-dependent *BCL6* suppression may take place in GC B cells undergoing terminal differentiation, correlating with reduced *BCL6* levels in plasma B cells [46]. MBD1 binding to *BCL6* promoter is observed in HCT116 cells (ChIP atlas:DRX021118 ;https://chip-atlas.org/view?id=DRX021118; Figure 2E). MBD1 binding may contribute to reduced genomic instability at the *BCL6* locus preventing *BCL6* rearregements as well as global rearrangements in GC B cells. On the other hand, shMBD1-Raji cells exhibited higher sensitivity to HU, gemcitabine, and etoposide as well as reduced tumorogencity in xenograft models than shSCR-Raji cells (Figure 5A). These results suggest that MBD1 promotes survival in cancer cells, making MBD1 an attractive therpautic target. The reduced tumorigenicity observed in the shMBD1-Raji xenograft could be due to the increased accumulation of genomic instability in shMBD1-Raji cells compared to shSCR-Raji cells (Figure 5C, D).

### AID mutation signatures in *MBD1*-mutant somatic cancers

AID-mediated mutagenesis drives B lymphomagenesis by mutating tumor suppresor genes and superenhancers, altering the binding of transcription factors and chromatin remodelers [10]. Interestingly, about 39% of single base-pair alterations in *MBD1* were C>T transition mutations (Figure 4F), a classic signature of AID-dependent mutagenesis (Figure 4F). Thus AID-dependent *MBD1* mutagenesis could serve as an important adapation for B lymphoma selection during the GC reaction.

### *MBD1* and *IRF4* expression in differentiating GC B cells and B-lymphomas

We found that *MBD1* and *IRF4* expression is positively correlated in DLBCL (Supplementary Figure 3A,B). On the other hand, *Mbd1* and *Irf4* were induced in mouse and human GC B cells undergoing plasma B differentiation (Figure 1C, Supplementary Figure 3C), indicating a synchronous upregulation of MBD1 and IRF4 in GC B cells undergoing differentiation. The mechanims of *MBD1* and *IRF4* co-expression in GC B cells is not yet clear. It is possible that *MBD1* and *IRF4* could be induced by decreased BCL6 levels in differetiating GC B cells. Also, MBD1 in differentiating GC B cells could suppreses *BCL6*, allowing for higher *IRF4* expression and plasma B cell differentiation. Importantly, *IRF4* suppression is mediated by EZH2-dependent H3K27me3 [23]. Is is possible that MBD1 may counteract EZH2 by recruiting histone deacetylases and transcriptional repressors to the *EZH2* promoter [5, 24, 33, 45]. This positions MBD1 as a novel regulator of GC B cell differentiation regulating the plasma B cell differentiation [47, 48]. Another possibility is that MBD1 may work alongside BCL6 and BACH2, both of which negatively regulate *IRF4* and *PRDM1* [46], allowing for homestasis of *IRF4* expression. This is consistent with our finding where *MBD1* depletion in Raji cells induced *IRF4* expression (Supplementary Figure 3D). However MBD1-mediated suppression of *IRF4* in Raji lymphoma cells may be distinct from WT B cells residing in the GC compartment, therefore we can not rule out the possibility that MBD1 promotes plasma B cell differentiation.

MBD1-mediated regulation of *IRF4* and *PRDM1* and its significance in the suppression and/or survival of B-lymphomas remains to be elucidated. We propose two potential regulatory scenarios for MBD1-dependent *IRF4* regulation depending on the stage of tumorigenesis. In the first scenario, GC B cells with high-affinity B cell receptors (BCR) might experience higher MBD1 expression, facilitating normal GC B cell differentiation. In the second, B lymphoma precursors— unproductive GC B cells with damaged BCRs—may experience reduced *MBD1* expression blocking the expression of plasma B cell differentiation genes including *IRF4* and mitigating the risk of malignant transformation into multiple myeloma (MM). Thus, MBD1 may differentially regulate plasma B cell differentiation in normal plasma B cells versus B lymphoma precursors.

In summary, our findings suggest that MBD1 is a key regulator of GC B cell differentiation, acting independently of both BCL6 and BACH2. Additionally, MBD1 may prevent BCL6 rearrangement in GC B cells, thereby inhibiting the formation of GCDBL. Finally, considering MBD1’s role in cell viability in cells treated with DNA-replication inhibitors, it could serve as a novel therapeutic target for relapsed and refractory B cell cancers that are resistant to chemotherapy.

## Declarations

### Ethical Approval

The use of human/animal samples (wherever applicable) was approved as per Institutional ethical board.

### Authors’ contributions

SKG designed the original hypothesis, performed experiments, and analyzed data. KO analyzed *Mbd1* transcripts in mouse plasma B cells and provided constructive feedback. SKG wrote the manuscript. JHB critically read the manuscript and provided constructive feedback. All authors read and agreed to manuscript.

### Availability of data and materials

All data needed to evaluate the conclusions in the paper are present in the main text and the supplementary materials.

## Material and Methods

### Human B lymphoma cultures, treatment with inhibitors and plasmid overexpression

Human B lymphoma Cell lines Raji were obtained from Department of Hematology, Kyoto university hospital, RIKEN cell bioresources, Japan. MCF7, MDA-MB-231, SUM149PT, MDA-MB-436 were available at the Cancer Research Institute, Kanazawa University. Raji and HEK293T cells were cultured in RPMI medium (30264-85, Nakalai) including the 10% fetal bovine serum (FBS), 1% Penicillin-Streptomycin & Amphotericin B solution (Waken# 161-23181) at 37°C maintaining the 5% CO_2_ concentration. Breast cancer cell lines MDA-MB-231, MDA-MB-436, MCF-7 cells were cultured DMEM, supplemented with 10% fetal bovine serum (FBS), 2 mM L-glutamine and 1 × Penicillin-Streptomycin (Life Technologies). SUM149PT cell were cultivated in Ham’s F-12 medium containing 5% FBS, 10 μg/ml insulin, 1% Penicillin-Streptomycin & Amphotericin B solution (Waken# 161-23181), and supplemented with 0.5 μg/ml hydrocortisone. Cells were propagated at 37 °C in a 5% CO_2_ atmosphere. FX1 (S8591, Selleck Japan), Gemcitabine (S1714, Selleck, Japan), Hydroxyurea (085-06653, Wako, Japan) and etoposide (055-0843, Wako, Japan) treatments were performed as described in figure legends. The preparation of these chemical was performed as per the manufacturer’s recommendation. For the overexpression of *pcDNA* and *pcDNA-BCL6* plasmids in Raji cells, the cells were cultured for 2 hours in RPMI medium containing 1% FBS, followed by transfection with Lipofectamine 3000 according to the manufacturer’s instructions (Thermo Fisher, Japan). Twelve hours post-transfection, the total FBS concentration was adjusted to 10% by adding additional FBS. Cells were collected for analysis at the indicated time points.

### shRNA preparation and lentivirus induced knockdown

HEK293T cells were cultured at the 70% confluency. For transfection mixture and lentivirus preparation, viral vector *PAX2*, *pVSVG* and *pLKO.1* plasmids (1 μg of each) cloned with respective shRNA sequences were mixed in 200 μl of warm Optimem medium followed by addition of 3 μg of PEI reagent. The mix incubated for 20 minutes at room temperature follwed by addition to the H3K293T cells. Fresh medium DMEM medium was replaced to transfected cells after 24 hours. The H3K293T cells were then cultured for additional 48 hours. The culture medium containing the lentivirus particles was then filtered using the 0.45-micron filters and the filtrate was aliquoted and stocked in -80 °C until used. For stable transfection of Raji cells, 1 million cells were transduced with 400 μl of filtered lentiviral supernatant follwed by selection with puromycin (2000 ng/ml) for 2 days. The 2 days selected cells were again washed and selected for additional 2 days with similar dose of Puromycin.

### RNA isolation, cDNA preparation, qPCR analysis and RNA-seq analysis

Total RNA was isolated with NucleoSpin® RNA Plus (#740984.50, Takara) and cDNA was prepared with the RT-PCR mix (FSQ#201, Toyobo Japan). qPCR was performed using the SYBR mix as per manufacturer’s instructions (Thunderbolt QPS-201, Toyobo Japan). Data were analyzed using the delta-delta Ct method indicating relative transcript levels to the respective control samples. ACTB gene signals were used as a housekeeping gene. Computational analyses of RNA-seq were performed as described (Reference; transcriptome paper). The RNA-seq and ChIP data set are under GEO accession numbers GSE242375 and GSE242936 respectively.

### Chromatin Immunoprecipitation (ChIP), ChIP-seq Library Preparation, Computational Analysis of ChIP-seq Data, and MBD1 Binding to *BCL6* locus withChIP-Atlas

ChIP was performed as previously described [6]. The ChIP-seq library was prepared as per the manufacturer’s instruction (Thruplex DNA-seq Kit, R400675, Takarabio, Japan). ChIP-seq was performed as described [31]. The ChIP-seq files were deposited to NCBI under accession number GSE242936. DNA CpG methylation status of mouse *Mbd1* promoter in B cells were reanalyzed using the wild type and *Sca1-Bcl6^Δ^* mice data [25, 43]. Binding of MDB1 on *BCL6* promoter was examined using the dataset # DRX021118 available on ChIP-atlas (https://chip-atlas.org/view?id=DRX021118).

### BCL6 binding motif, luciferase assay and COSMIC datasets

The luciferase plasmids depleting the BCL6 binding motifs on the *MBD1* promoter were constructed using the Q5 site-directed mutagenesis kit (NEB# E0554S) and as per the manufacturer’s instructions. Primers used for the site-directed mutagenesis are listed in Table 1. The plasmids were transfected in the HEK293T cells for 24 hours using lipofectamine-3000 (ThermoFisher# L3000001). After 24 hours, cell lysates were prepared using Promega luciferase assay kit. The Catalogue of somatic mutation in cancers (COSMIC) datasets were searched for cancers with *MBD1* mutation and the mutation signatures were classified and filtered for the AID signatures.

### Tumor xenograft in nude mouse and estimation of cell viability

For tumor xenograft, and injection of Raji into nude mice, 0.6 million cells were suspended in 200 μl of Matrigel and medium mix, suspended well and subcutaneously injected into the right arm of the nude mice. The tumor growth was observed for 55 days and mice were sacrificed, followed by isolation of tumor and weighing. Treatment with chemotherapeutics and cell viability was tested by the cell viability kit (Dojindo Japan) as per the manufacturer’s instructions.

**Supplementary Table 1.**
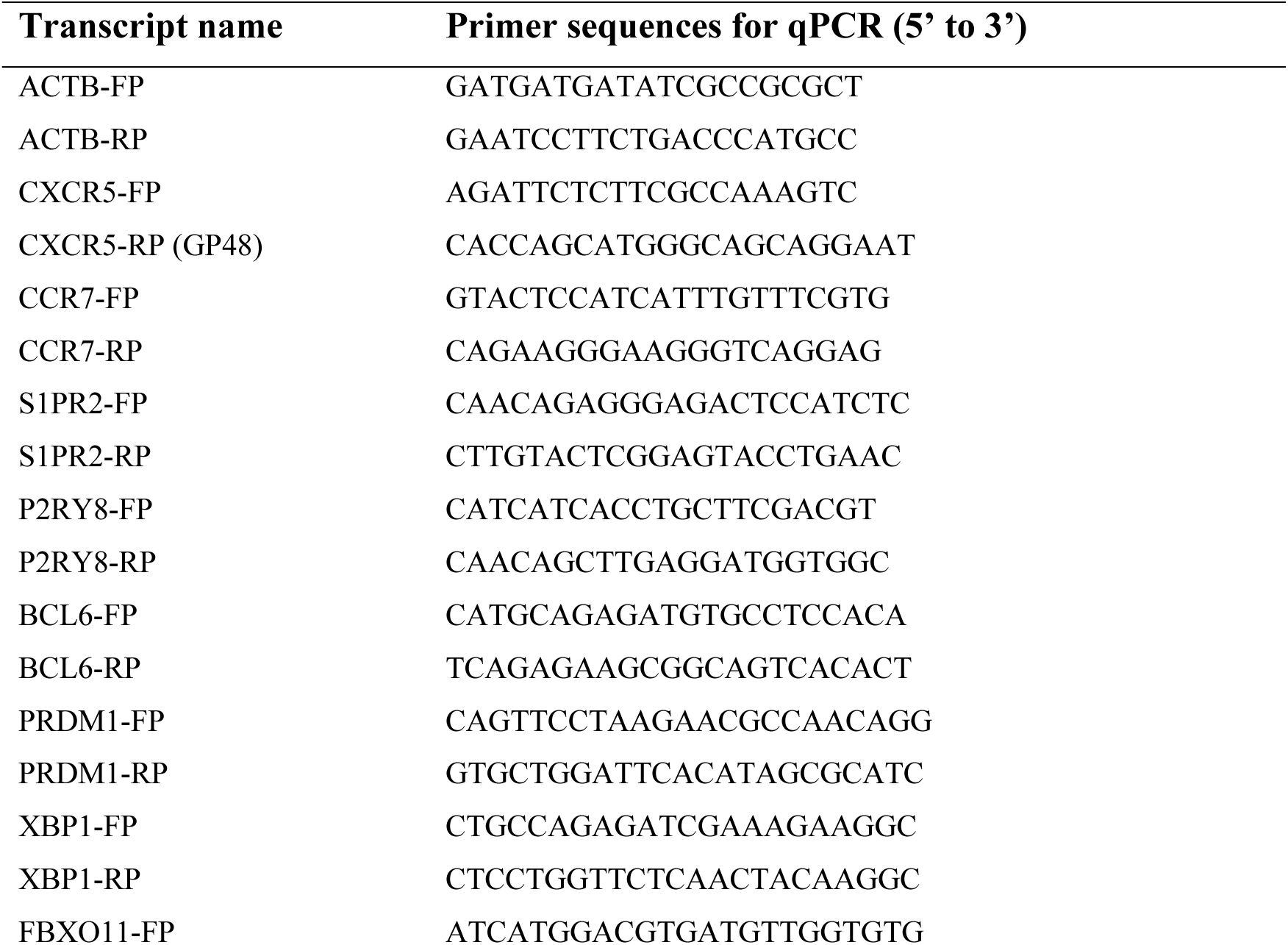
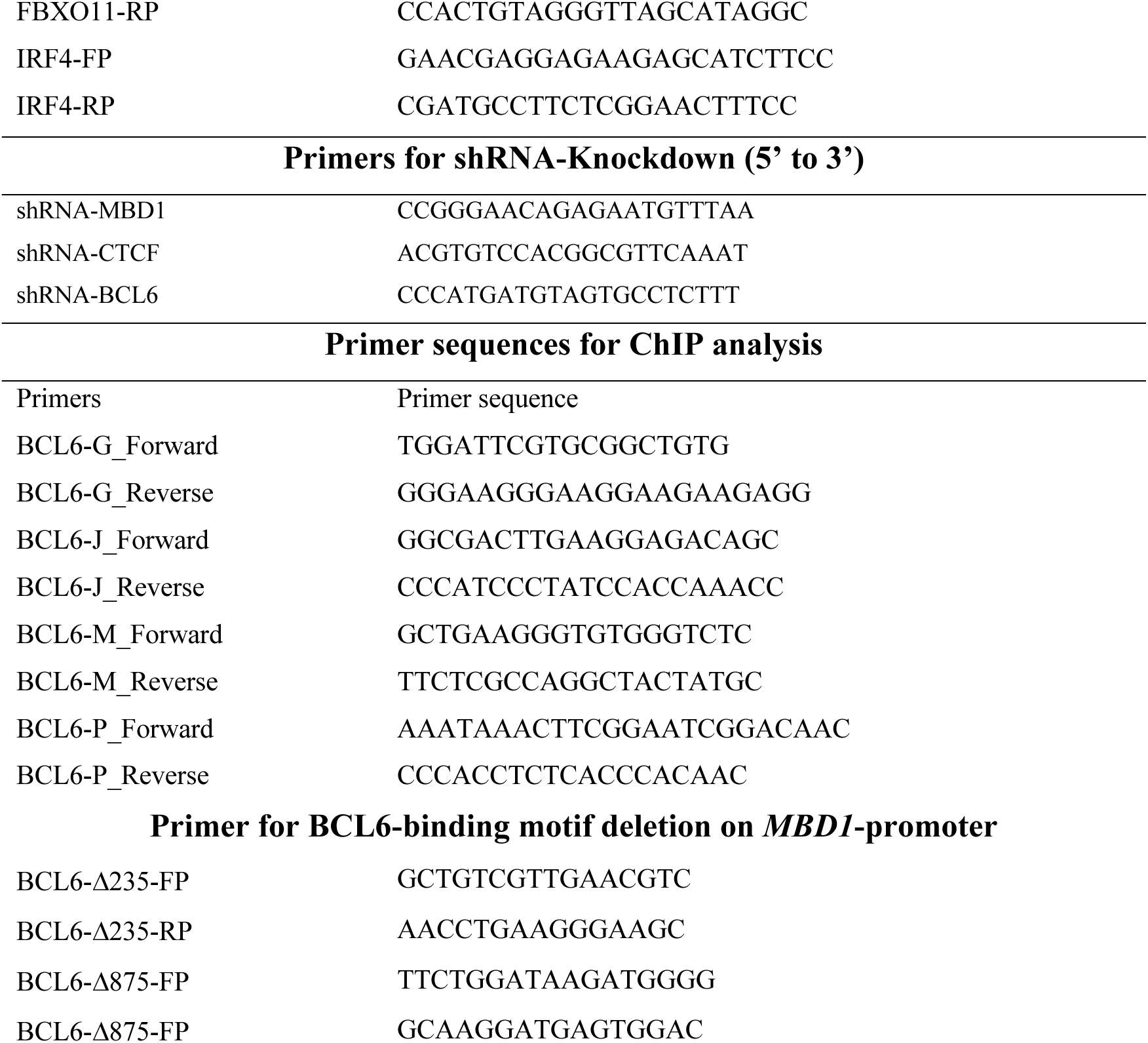
for primer sequences.

**Supplementary Table-2.**

| Antibody name | Purpose | Product/Manufacturer |
| --- | --- | --- |
| BCL6 | WB | GeneTex#GTC101338 |
| Histone H3K4-trimethyl | ChIP | CST#C42D8 |
| ACTB | WB | Novusbio (NB100-56874) |
| Histone H3 | WB | Biolegend (819411) |
| γH2AX | ChIP | Abcam#ab20669 |
| CTCF | ChIP, WB, IP | Active Motif#61932 |
| MBD1 | WB | Abeomics (ABM15H2) |

**Supplementary Table 3.** List of tumor acronym from TCGA study datasets.

| Study Abbreviation | Study Name |
| --- | --- |
| LAML | Acute Myeloid Leukemia |
| ACC | Adrenocortical carcinoma |
| BLCA | Bladder Urothelial Carcinoma |
| BRCA | Breast invasive carcinoma |
| CESC | Cervical squamous cell carcinoma and endocervical adenocarcinoma |
| CHOL | Cholangiocarcinoma |
| COAD | Colon adenocarcinoma |
| ESCA | Esophageal carcinoma |
| GBM | Glioblastoma multiforme |
| HNSC | Head and Neck squamous cell carcinoma |
| KICH | Kidney Chromophobe |
| KIRC | Kidney renal clear cell carcinoma |
| KIRP | Kidney renal papillary cell carcinoma |
| LIHC | Liver hepatocellular carcinoma |
| LUAD | Lung adenocarcinoma |
| LUSC | Lung squamous cell carcinoma |
| DLBC | Lymphoid Neoplasm Diffuse Large B cell Lymphoma |
| PAAD | Pancreatic adenocarcinoma |
| PCPG | Pheochromocytoma and Paraganglioma |
| PRAD | Prostate adenocarcinoma |
| READ | Rectum adenocarcinoma |
| SKCM | Skin Cutaneous Melanoma |
| STAD | Stomach adenocarcinoma |
| TGCT | Testicular Germ Cell Tumors |
| THYM | Thymoma |
| UCS | Uterine Carcinosarcoma |
| UCEC | Uterine Corpus Endometrial Carcinoma |
| UVM | Uveal Melanoma |

**Supplementary Figure 1.**
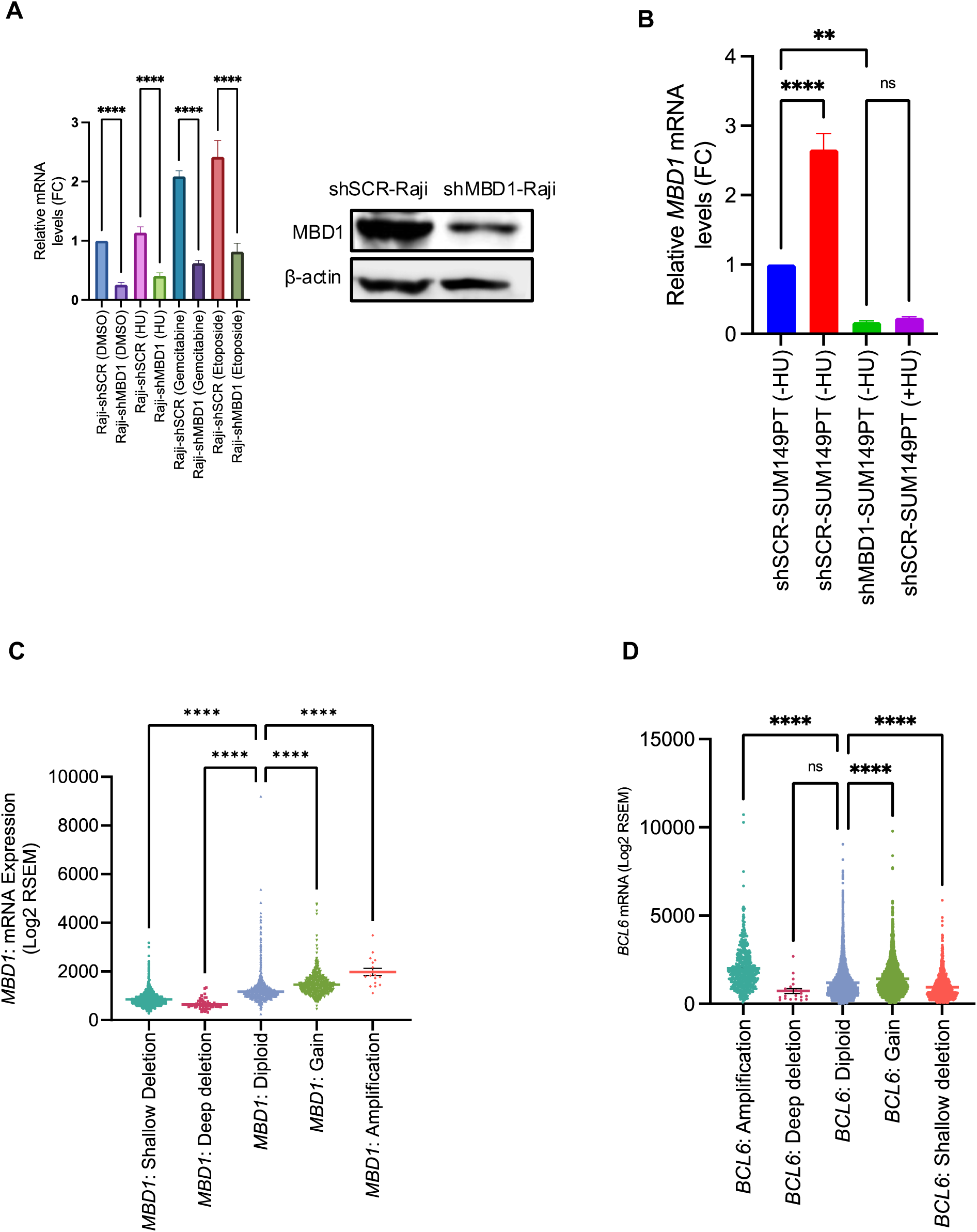
**(A)** Quantitative PCR (qPCR) analysis of *MBD1* mRNA levels in shSCR-Raji and shMBD1-Raji cells treated with DMSO, hydroxyurea (HU, 10 mM), gemcitabine (20 mM), and etoposide (50 μM) for 24 hours. Data are presented as mean ± SEM, n = 3. Tukey’s multiple comparison test results: *p < 0.0001* for shSCR-Raji (DMSO) vs. shMBD1-Raji (DMSO) groups, *p < 0.0001* for shSCR-Raji (DMSO) vs. shSCR-Raji (gemcitabine) groups, p < 0.0001 for shSCR-Raji (DMSO) vs. shSCR-Raji (etoposide) groups, p = 0.8068 for shMBD1-Raji (DMSO) vs. shMBD1-Raji (HU) groups, *p=0.0398* for shMBD1-Raji (DMSO) vs. shMBD1-Raji (gemcitabine) groups, and *p < 0.0010* for shMBD1-Raji (DMSO) vs. shMBD1-Raji (etoposide) groups. Western blot image of MBD1 using cell lysates from shSCR-Raji and shMBD1-Raji cells **(B)** qPCR analysis of MBD1 mRNA in shSCR-SUM149PT and shMBD1-SUM149PT cells treated with or without HU (10 mM) for 24 hours. Tukey’s multiple comparison test: *p < 0.001* for shSCR-SUM149PT (-HU) vs. shSCR-SUM149PT (+10 mM HU) cells **(C)** *MBD1* mRNA expression profile among the TCGA tumor samples. Mean *MBD1* mRNA RSEM (log2+1) value was 850.3 in *MBD1*-shallow deletion (n=3208) , 646 in *MBD1*-deletions (n=56), 1171 in *MBD1*-diploid (n=5789), 1465 in *MBD1*-gain (n=819), 1979 in *MBD1*-amplification (n=17) groups respectively. *p< 0.0001* for *MBD1*-diploid vs *MBD1*-amplification, *p< 0.0001* for *MBD1*-diploid vs *MBD1* Deep-deletion, *p< 0.0001* for *MBD1*-diploid vs *MBD1*-gain, *p< 0.0001* for *MBD1*-diploid vs *MBD1*-shallow deletion. Dunnett’s multiple comparison test, One way ANOVA. **(D)** *BCL6* mRNA expression profile among the TCGA tumor samples. Mean *BCL6* mRNA value (RSEM:log2+1) was 2006 for *BCL6*-amplification (n=488), 726.3 for *BCL6*-deep deletions (n=21), 1198 for *BCL6*-diploid (n=5950), 1417 for *BCL6*-gain (n=2587) and 949.6 for *BCL6*-shallow deletion (n=864) group. *p< 0.0001* for *BCL6*-diploid vs *BCL6*-amplification, *p= 0.08* for *BCL6*-diploid vs *BCL6* Deep-deletion, *p< 0.0001* for *BCL6*-diploid vs *BCL6*-gain, *p< 0.0001* for *BCL6*-diploid vs *BCL6*-shallow deletion. Dunnett’s multiple comparison test, One way ANOVA.

**Supplementary Figure 2.**
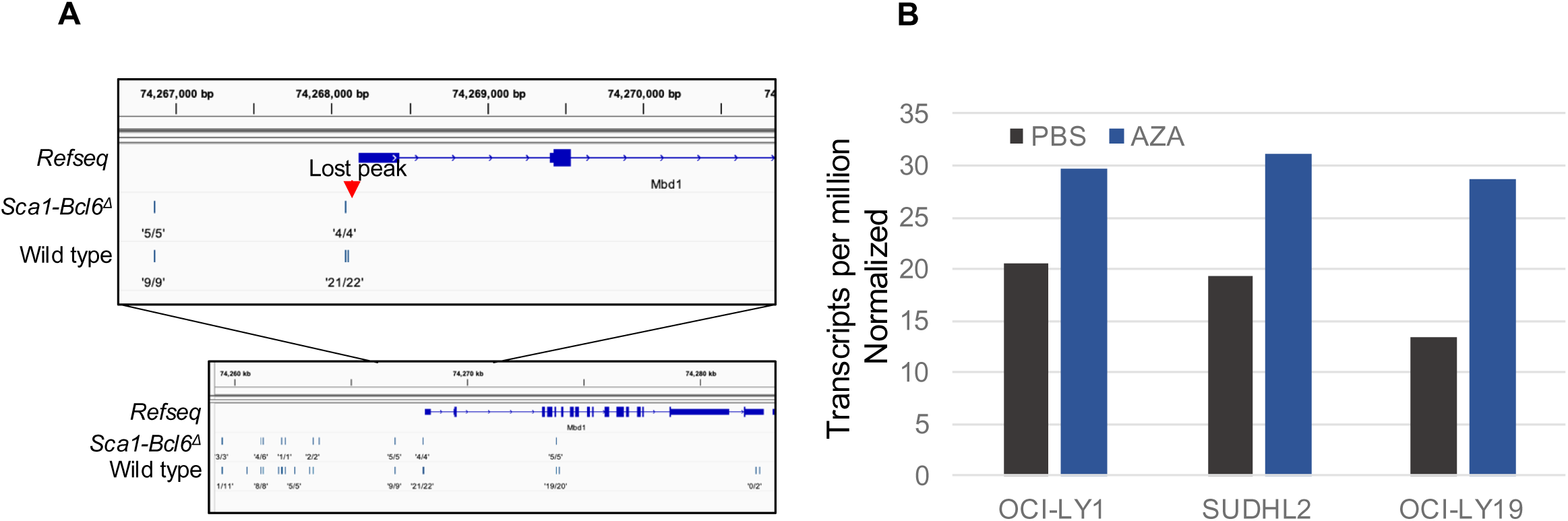
*MBD1* expression is associated with DNA methylation and *Bcl6* status in mouse GC B cells. **(A)** CpG methylation status of activated mouse GC B cell in wild type and *Sca1-Bcl6^Δ^* mice were analyzed from published study [25, 43]. The CpG methylation signal on position 4’4’ of wild type *Mbd1* locus are lost in *Sca1-Bcl6^Δ^* **(B)** AZA treatment of OCI-LY1, SUDHL2 and OCI-LY19 induced *MBD1* mRNA expression (GSE190319). Y-axis indicated values of *MBD1* mRNA in counts of Transcription per million.

**Supplementary Figure 3.**
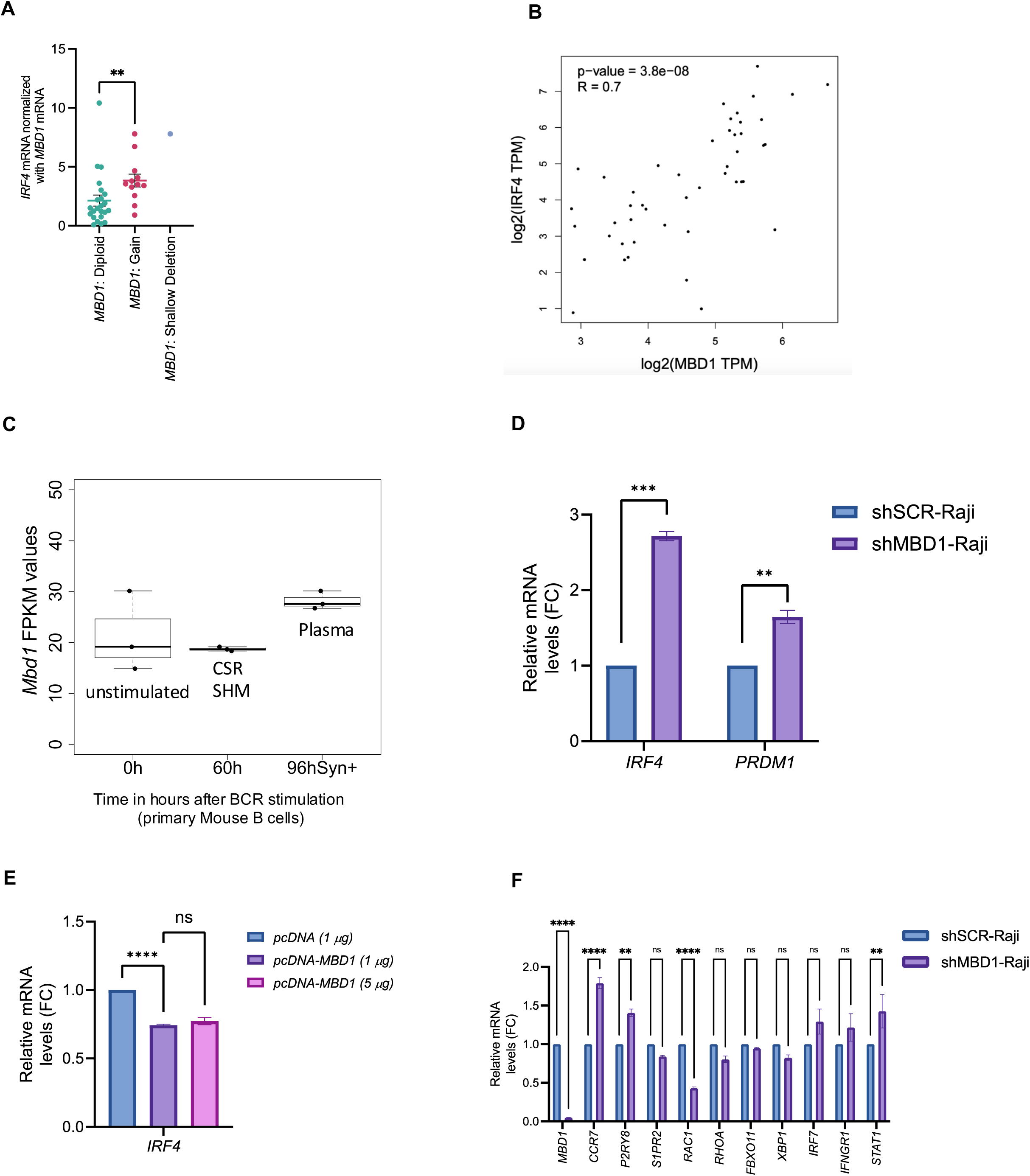
Correlation of *MBD1* and *IRF4* expression in DLBCL tumors available at TCGA database. DLBCL tumors exhibiting *BACH2*-diploid and *IRF4-*diploid status were sorted and catergorised into *MBD1*-diploid (n=23), *MBD1*-gain (n=12) and *MBD1*-shallow deletion (n=1) groups. The Y-axis displays normalized values of *IRF4* mRNA (RSEM counts) with the *MBD1* mRNA (RSEM counts). p=0.0051 for *MBD1*-diploid vs *MBD1*-gain tumors. Mann Whiteny t-test. **(B)** Correlation of *MBD1* and *IRF4* co-expression in DLBCL samples obtained from GEPIA. R=0.7, p=3.8e-08. **(C)** *Mbd1* expression is induced in activated mouse germinal center B cells at the stage of plasma cell differentiation (96 hours) compared to unstimulated (0 hours) and germinal center B cells undergoing peak class switch recombination (CSR) and somatic hypermutation (SHM). **(D)** qPCR analysis of *IRF4*, and *PRDM1* in shSCR-Raji and shMBD1-Raji cells. Data are presented as mean ± SEM, n = 3. Unpaired t-test. For *IRF4*, *p <0.0001* between shSCR-Raji vs. shMBD1-Raji, for *PRDM1* p =0.0018 between shSCR-Raji vs. shMBD1-Raji cells **(E)** Raji cells were transfected with either the control *pcDNA* vector or the *pcDNA-MBD1* plasmid for 24 hours, followed by quantification of *IRF4* mRNA levels using qRT-PCR. Unpaired t-test. p <0.0001 for *pcDNA* (1 μg) vs. *pcDNA-MBD1* (1 μg); p = 0.3360 for *pcDNA-MBD1* (1 μg) vs. *pcDNA-MBD1* (5 μg). Data are presented as mean ± SEM, n=3 **(F)** qPCR analysis of GC marker genes in shSCR-Raji and shMBD1-Raji cells. *MBD1*; p <0.0001 for shSCR-Raji vs shMBD1-Raji cells. *CCR7*; p <0.0001 for shSCR-Raji vs shMBD1-Raji cells. *P2RY8*; p =0.0030 for shSCR-Raji vs shMBD1-Raji cells. *S1PR2*; p =0.7833 for shSCR-Raji vs shMBD1-Raji cells. *RAC1*; p <0.0001 for shSCR-Raji vs shMBD1-Raji cells. *RHOA*; p =0.5067 for shSCR-Raji vs shMBD1-Raji cells. *FBXO11*; p >0.9999 for shSCR-Raji vs shMBD1-Raji cells. *XBP1*; p =0.6600 for shSCR-Raji vs shMBD1-Raji cells. *IRF7*; p =0.0724 for shSCR-Raji vs shMBD1-Raji cells. *IFNGR1*; p =0.3715 for shSCR-Raji vs shMBD1-Raji cells. *STAT1*; p =0.0017 for shSCR-Raji vs shMBD1-Raji cells. Sidak’s multiple comparisons test. Data are presented as mean ± SEM, n=3.

## References

1. Allen, C.D., et al., Germinal center dark and light zone organization is mediated by CXCR4 and CXCR5. Nat Immunol, 2004. 5(9): p. 943–52.

2. Victora, G.D. and M.C. Nussenzweig, Germinal Centers. Annu Rev Immunol, 2022. 40: p. 413–442.

3. Muramatsu, M., et al., Class switch recombination and hypermutation require activation-induced cytidine deaminase (AID), a potential RNA editing enzyme. Cell, 2000. 102(5): p. 553–63.

4. Roco, J.A., et al., Class-Switch Recombination Occurs Infrequently in Germinal Centers. Immunity, 2019. 51(2): p. 337–350.e7.

5. Lu, Z., et al., BCL6 breaks occur at different AID sequence motifs in Ig-BCL6 and non-Ig-BCL6 rearrangements. Blood, 2013. 121(22): p. 4551–4.

6. Gothwal, S.K., et al., BRD2 promotes antibody class switch recombination by facilitating DNA repair in collaboration with NIPBL. Nucleic Acids Res, 2024.

7. Shaffer, A.L., et al., IRF4 addiction in multiple myeloma. Nature, 2008. 454(7201): p. 226–31.

8. Kaiser, L.M., et al., CXCR4 in Waldenström’s Macroglobulinema: chances and challenges. Leukemia, 2021. 35(2): p. 333–345.

9. Kurtova, A.V., et al., Mantle cell lymphoma cells express high levels of CXCR4, CXCR5, and VLA-4 (CD49d): importance for interactions with the stromal microenvironment and specific targeting. Blood, 2009. 113(19): p. 4604–13.

10. Bal, E., et al., Super-enhancer hypermutation alters oncogene expression in B cell lymphoma. Nature, 2022. 607(7920): p. 808–815.

11. Calpe, E., et al., ZAP-70 promotes the infiltration of malignant B-lymphocytes into the bone marrow by enhancing signaling and migration after CXCR4 stimulation. PLoS One, 2013. 8(12): p. e81221.

12. Dhordain, P., et al., The LAZ3(BCL-6) oncoprotein recruits a SMRT/mSIN3A/histone deacetylase containing complex to mediate transcriptional repression. Nucleic Acids Res, 1998. 26(20): p. 4645–51.

13. Chang, C.C., et al., BCL-6, a POZ/zinc-finger protein, is a sequence-specific transcriptional repressor. Proc Natl Acad Sci U S A, 1996. 93(14): p. 6947–52.

14. Phan, R.T., et al., BCL6 interacts with the transcription factor Miz-1 to suppress the cyclin-dependent kinase inhibitor p21 and cell cycle arrest in germinal center B cells. Nat Immunol, 2005. 6(10): p. 1054–60.

15. Akkaya, M., K. Kwak, and S.K. Pierce, B cell memory: building two walls of protection against pathogens. Nature Reviews Immunology, 2020. 20(4): p. 229–238.

16. Ye, B.H., et al., Alterations of a zinc finger-encoding gene, BCL-6, in diffuse large-cell lymphoma. Science, 1993. 262(5134): p. 747–50.

17. Ci, W., J.M. Polo, and A. Melnick, B-cell lymphoma 6 and the molecular pathogenesis of diffuse large B-cell lymphoma. Curr Opin Hematol, 2008. 15(4): p. 381–90.

18. Ruggieri, S., et al., Translocation of the proto-oncogene Bcl-6 in human glioblastoma multiforme. Cancer Letters, 2014. 353(1): p. 41–51.

19. Duy, C., et al., BCL6 enables Ph+ acute lymphoblastic leukaemia cells to survive BCR-ABL1 kinase inhibition. Nature, 2011. 473(7347): p. 384–8.

20. Walker, S.R., et al., The transcriptional modulator BCL6 as a molecular target for breast cancer therapy. Oncogene, 2015. 34(9): p. 1073–82.

21. Xu, L., et al., BCL6 promotes glioma and serves as a therapeutic target. Proc Natl Acad Sci U S A, 2017. 114(15): p. 3981–3986.

22. Shaknovich, R., et al., DNA methyltransferase 1 and DNA methylation patterning contribute to germinal center B-cell differentiation. Blood, 2011. 118(13): p. 3559–69.

23. Béguelin, W., et al., EZH2 is required for germinal center formation and somatic EZH2 mutations promote lymphoid transformation. Cancer Cell, 2013. 23(5): p. 677–92.

24. Villa, R., et al., The methyl-CpG binding protein MBD1 is required for PML-RARalpha function. Proc Natl Acad Sci U S A, 2006. 103(5): p. 1400–5.

25. Caron, G., et al., Cell-Cycle-Dependent Reconfiguration of the DNA Methylome during Terminal Differentiation of Human B Cells into Plasma Cells. Cell Reports, 2015. 13(5): p. 1059–1071.

26. Lai, A.Y., et al., DNA methylation prevents CTCF-mediated silencing of the oncogene BCL6 in B cell lymphomas. Journal of Experimental Medicine, 2010. 207(9): p. 1939–1950.

27. Bell, A.C. and G. Felsenfeld, Methylation of a CTCF-dependent boundary controls imprinted expression of the Igf2 gene. Nature, 2000. 405(6785): p. 482–5.

28. Hark, A.T., et al., CTCF mediates methylation-sensitive enhancer-blocking activity at the H19/Igf2 locus. Nature, 2000. 405(6785): p. 486–9.

29. Kosak, S.T., et al., Subnuclear compartmentalization of immunoglobulin loci during lymphocyte development. Science, 2002. 296(5565): p. 158–62.

30. Jhunjhunwala, S., et al., The 3D structure of the immunoglobulin heavy-chain locus: implications for long-range genomic interactions. Cell, 2008. 133(2): p. 265–79.

31. Gothwal, S.K. and J.H. Barlow, Stress Induced Signaling Pathways in Burkitt Lymphoma Play Novel Mechanisms in Fate Determination and Pathogenesis of Germinal Center-Derived B-Lymphomas. bioRxiv, 2024: p. 2024.12.19.628635.

32. Holmes, A.B., et al., Single-cell analysis of germinal-center B cells informs on lymphoma cell of origin and outcome. Journal of Experimental Medicine, 2020. 217(10).

33. Fujita, N., et al., Methyl-CpG binding domain 1 (MBD1) interacts with the Suv39h1-HP1 heterochromatic complex for DNA methylation-based transcriptional repression. J Biol Chem, 2003. 278(26): p. 24132–8.

34. Clouaire, T., et al., Recruitment of MBD1 to target genes requires sequence-specific interaction of the MBD domain with methylated DNA. Nucleic Acids Res, 2010. 38(14): p. 4620–34.

35. Ci, W., et al., The BCL6 transcriptional program features repression of multiple oncogenes in primary B cells and is deregulated in DLBCL. Blood, 2009. 113(22): p. 5536–48.

36. Cardenas, M.G., et al., Rationally designed BCL6 inhibitors target activated B cell diffuse large B cell lymphoma. J Clin Invest, 2016. 126(9): p. 3351–62.

37. Saito, M., et al., A signaling pathway mediating downregulation of BCL6 in germinal center B cells is blocked by BCL6 gene alterations in B cell lymphoma. Cancer Cell, 2007. 12(3): p. 280–92.

38. Green, M.R., et al., Transient expression of Bcl6 is sufficient for oncogenic function and induction of mature B-cell lymphoma. Nature Communications, 2014. 5(1): p. 3904.

39. Oprea, M. and A.S. Perelson, Somatic mutation leads to efficient affinity maturation when centrocytes recycle back to centroblasts. J Immunol, 1997. 158(11): p. 5155–62.

40. Ochiai, K., et al., Accelerated plasma-cell differentiation in Bach2-deficient mouse B cells is caused by altered IRF4 functions. The EMBO Journal, 2024. 43(10): p. 1947–1964-1964.

41. Margiotta, A. and C. Bucci, Coordination between Rac1 and Rab Proteins: Functional Implications in Health and Disease. Cells, 2019. 8(5).

42. Green, J.A. and J.G. Cyster, S1PR2 links germinal center confinement and growth regulation. Immunol Rev, 2012. 247(1): p. 36–51.

43. Kulis, M., et al., Whole-genome fingerprint of the DNA methylome during human B cell differentiation. Nat Genet, 2015. 47(7): p. 746–56.

44. Amanda, S., et al., IRF4 drives clonal evolution and lineage choice in a zebrafish model of T-cell lymphoma. Nat Commun, 2022. 13(1): p. 2420.

45. Fujita, N., et al., Mechanism of transcriptional regulation by methyl-CpG binding protein MBD1. Mol Cell Biol, 2000. 20(14): p. 5107–18.

46. Huang, C., et al., Cooperative transcriptional repression by BCL6 and BACH2 in germinal center B-cell differentiation. Blood, 2014. 123(7): p. 1012–20.

47. Kuo, T.C., et al., Repression of BCL-6 is required for the formation of human memory B cells in vitro. J Exp Med, 2007. 204(4): p. 819–30.

48. Lin, K.I., C. Tunyaplin, and K. Calame, Transcriptional regulatory cascades controlling plasma cell differentiation. Immunol Rev, 2003. 194: p. 19–28.

